# Recovery of the full *in vivo* firing range in post-lesion surviving DA SN neurons associated with Kv4.3-mediated pacemaker plasticity

**DOI:** 10.1101/2021.08.09.455657

**Authors:** Lora Kovacheva, Josef Shin, Josefa Zaldivar-Diez, Johanna Mankel, Navid Farassat, Kauê Machado Costa, Poonam Thakur, José Obeso, Jochen Roeper

**Affiliations:** Institute of Neurophysiology, Neuroscience Center, Goethe University, Frankfurt, Germany; Department of Neurology, University Medical Centre of the Johannes Gutenberg University Mainz, Mainz, Germany; HM CINAC (Centro Integral de Neurociencias Abarca Campal), Hospital Universitario HM Puerta del Sur, HM Hospitales. Madrid, Spain. CIBERNED, Instituto de Salud Carlos III, Madrid, Spain. Medical School, CEU-San Pablo University, Madrid, Spain; Eye Center, Medical Center-University of Freiburg, Freiburg im Breisgau, Germany; Department of Psychology, University of Alabama at Birmingham, AL, USA; School of Biology, Indian Institute of Science Education and Research (IISER)-Thiruvananthapuram, Kerala, India

**Keywords:** dopamine, 6-OHDA, Kv4.3 potassium channel, substantia nigra, pacemaker, homeostasis, intrinsic plasticity

## Abstract

Dopamine (DA) neurons in the substantia nigra (SN) control several essential functions, including the voluntary movement, learning and motivated behavior. Healthy DA SN neurons show diverse firing patterns *in vivo*, ranging from slow pacemaker-like activity (1-10 Hz) to transient high frequency bursts (<100 Hz), interspersed with pauses that can last hundreds of milliseconds. Recent *in vivo* patch experiments have started to reveal the subthreshold mechanisms underlying this physiological diversity, but the impact of challenges like cell loss on the *in vivo* activity of adult DA SN neurons, and how these may relate to behavioral disturbances, are still largely unknown. We investigated the *in vivo* electrophysiological properties of surviving SN DA neurons after partial unilateral 6-OHDA lesions, a single-hit, non-progressive model of neuronal cell loss. We show that mice subjected to this model have an initial motor impairment, measured by asymmetrical rotations in the open field test, which recovered over time. At 3 weeks post-lesion, when open field locomotion was strongly impaired, surviving DA SN neurons showed a compressed *in vivo* dynamic firing range, characterized by a 10-fold reduction of *in vivo* burst firing compared to controls. This *in vivo* phenotype was accompanied by pronounced *in vitro* pacemaker instability. In contrast, in the chronic post-lesion phase (>2 months), where turning symmetry in open field locomotion had recovered, surviving SN DA neurons displayed the full dynamic range of *in vivo* firing, including *in vivo* bursting, similar to controls. The normalized *in vivo* firing pattern was associated with a 2-fold acceleration of stable *in vitro* pacemaking, mediated by Kv4.3 potassium channel downregulation. Our findings demonstrate the existence of a homeostatic pacemaker plasticity mechanism in surviving DA SN neurons after pronounced cell loss.

## Introduction

Midbrain dopamine (DA) neurons in the substantia nigra (SN) project mainly to the dorsal striatum (DS), where they release dopamine in an activity-dependent manner, further shaped by additional mechanisms at the distal axon (Liu et al., 2022, Kramer et al., 2023). The resulting temporal and spatial dynamics of DA release in DS are relevant for multiple functions, including the control of voluntary movements, as evidenced by the cardinal features of Parkinson disease (PD), which is characterized by the death of SN DA neurons. The *in vivo* activity pattern of healthy SN DA neurons is complex, ranging from slow pacemaker-like activity (1-10 Hz) to transient high frequency bursts (<100 Hz), and interspersed with pauses that can last hundreds of milliseconds. However, our understanding if and how SN DA neuron electrical activity adapts to cell loss, relevant during aging or PD, is limited.

We have previously shown that overexpression of mutant A53T-α-synuclein induced an age-dependent acceleration of *in vivo* and *in vitro* firing frequencies. This was associated with redox-mediated dysfunction of Kv4.3 channels and an upregulation of Kv4.3 expression, potentially representing a homeostatic attempt to regulate pacemaker frequency (Subramaniam, Althof, et al. 2014). This study, however, left a more fundamental question unanswered: do adult DA SN neurons have the ability for homeostatic regulation of firing pattern in response to challenge? For this purpose, we employed the well-established 6-Hydroxy-Dopamine (6-OHDA) chemical lesion model in order to induce an unilateral loss of about half of the DA SN neurons. 6-OHDA has been widely used for generating non-progressive loss of DA SN neurons (Rubi and Fritschy 2020; P. R. L. Parker, Lalive, and Kreitzer 2016; Ungerstedt 1968). We opted for an unilateral, partial, intrastriatal 6-OHDA model, previously well characterized by Bez et al. (Bez, Francardo, and Cenci 2016). Bez and colleagues, as well as others, demonstrated that these partial 6-OHDA DA lesions are non-progressive and follow a stereotypical time course: an initial damage phase dominated by cell loss, inflammation and behavioral impairment for about 3 weeks, which is followed by a chronic phase (studied for up to 20 month in mice), characterized by partial behavioral recovery, neurochemical and molecular adaptations, and – to a certain degree – axonal sprouting of remaining DA neurons (Bez, Francardo, and Cenci 2016; Cenci and Björklund 2020; Kirik, Rosenblad, and Björklund 1998; Schwarting and Huston 1996; Winkler et al. 2002). This framework, with its distinct phases of impairment and subsequent partial functional recovery, provides a well-suited platform to investigate if and how the *in vivo* firing properties of surviving DA SN neurons change over time. Previous electrophysiological investigations of viable post-6-OHDA DA neurons have been limited, with only Hollerman & Grace providing a pioneering dataset (Hollerman and Grace 1990). They found lesion-size-dependent changes in *in vivo* firing properties of putative DA neurons in rats, including a decrease of *in vivo* burst firing occurring with larger lesions (Hollerman and Grace 1990). Therefore, conducting electrophysiological investigations at two distinct time-points within the context of a non-progressive partial 6-OHDA lesion is expected to provide novel insights into the homeostatic properties of identified DA SN neurons.

Here, we explore the electrophysiological properties, both *in vivo* and *in vitro,* of surviving SN DA neurons in an intra-striatal unilateral 6-OHDA lesion mouse model, titrated to about 40% surviving SN DA neurons. We selected two time points for recordings based on the continuous behavioral characterization of the motor phenotypes.

## Results

### Early behavioral impairments and long-term adaptations after partial lesion of substantia nigra dopamine neurons

To characterize electrophysiological properties in surviving and identified SN DA neurons, we employed a C57Bl6N mouse model with a partial 6-OHDA lesion of the nigrostriatal pathway. We explored different 6-OHDA concentrations (0.2µg/µl – 2µg/µl) and volumes (2µL and 6µL) resulting in a range of 0.4µg to 12µg 6-OHDA) to titrate the ipsilateral loss of DA SN neurons (Suppl. Fig. 1E for details). To target 40-50% cell loss, we slowly (250 nl/min) infused 6 µl of 2µg/µl 6-OHDA (i.e. 24 min total infusion time, total of 12 µg 6-OHDA) into the right DS. For controls, we infused vehicle (ACSF) with the same rate and same volume into the right DS without observing any damage (Fig. 1B; Suppl. Fig. 1-C). The loss of TH-positive SN neurons was quantified by unbiased stereology at 21 days and 3 months post-lesion (Fig. 1A).

**Figure 1.**
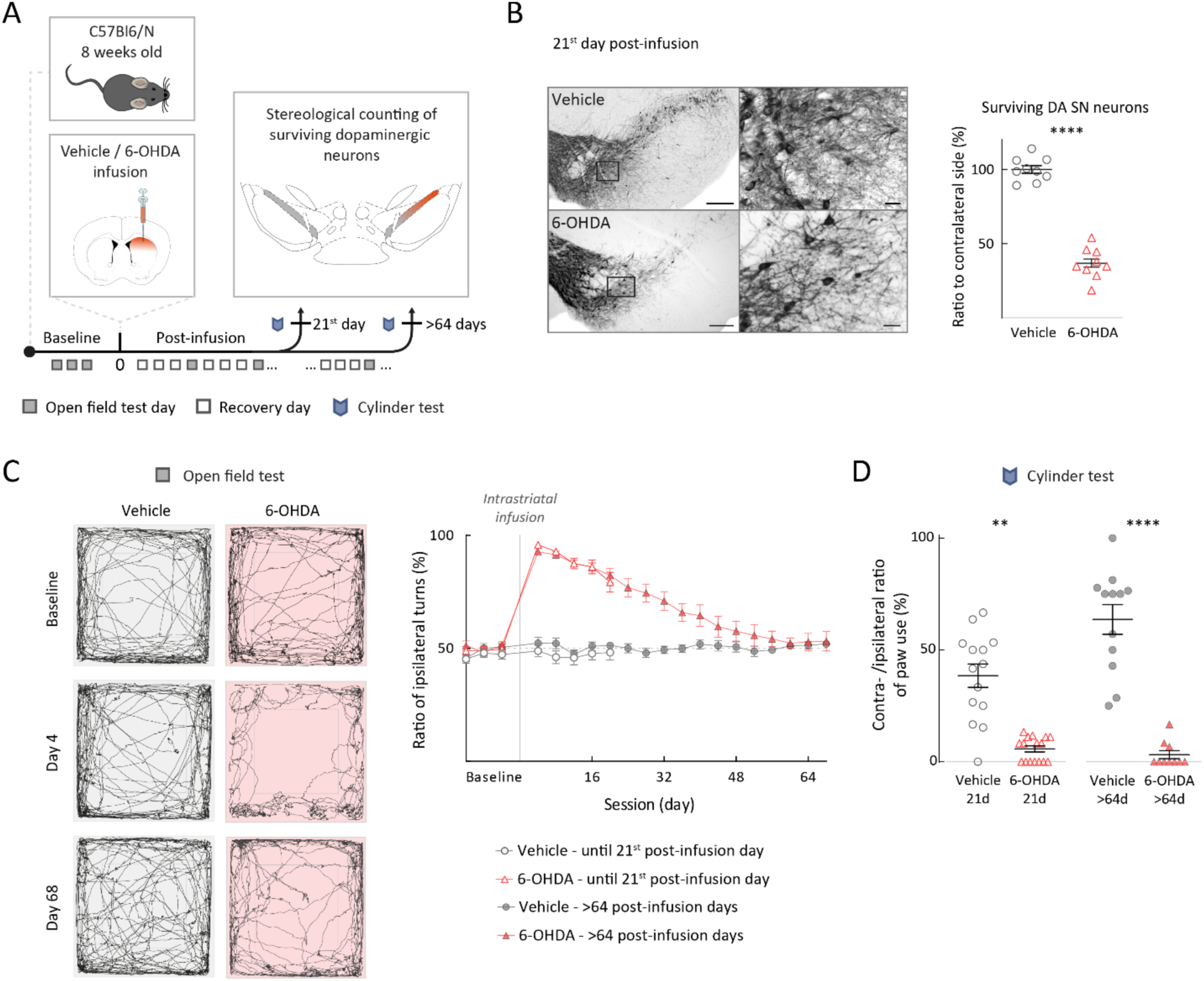
Unilateral striatal 6-OHDA mouse model with stable loss of SN DA neurons resulted in delayed partial behavioral recovery. (A) Experimental design, illustrating timeline of behavioral assays with termination points – groups ended either on the 21^st^, or later than the 64^th^ post-infusion day. (B) Left panel: TH-DAB staining of the SN, 10x magnification and 60x magnification for vehicle and 6-OHDA-infused mouse. Scalebar left 200µm, scalebar right 25µm. Right panel: ratio of ipsilateral (infusion side) to contralateral side of surviving TH-positive neurons in SN at 21^st^ post-infusion day. (C) Left: spontaneous locomotion of mice in open field arena for a 10 min session. Left, examples of an ACSF-infused mouse (vehicle), right example of a 6-OHDA-infused mouse at baseline (upper panels), 4^th^ post-infusion day (middle panels) and 68^th^ post-infusion day (lower panels). Right: ratio of ipsilateral to contralateral turning behavior for all experimental groups plotted against session days. Note recovery in the 6-OHDA-treated mice from day 4 to day 68 after initial strong turning bias (>90%). (D) Cylinder test quantified by ratio of contra-to ipsilateral forepaw use. Note significant loss of contralateral forepaw involvement, both at 21 and 64 days. All data are presented as mean ± standard error of mean (SEM).

We continuously characterized the motor phenotype of the model (i.e. open field locomotion, cylinder test) until the two different end points, either 21-days post-lesion (early phase) or more than 60 days post-lesion (late phase). In the first cohorts, the mice were sacrificed for TH immunohistochemistry in midbrain and striatum. As shown in Fig. 1B, at the early phase, the number of TH-positive (i.e. dopaminergic) SN neurons was reduced post-6-OHDA (lower left panels) compared to vehicle-control mice (upper left panels). The right panel in Fig. 1B displays the stereological results of TH-positive SN neurons for N=9 vehicle-treated and N=9 6-OHDA-infused male mice 21 days after their respective infusions. The data were normalized to the stereological results of respective contralateral SN DA neurons. In contrast to vehicle-infusions, where no ipsilateral loss of SN DA neurons was detected, the number of ipsilateral surviving SN DA neurons in the early post-6-OHDA phase was on average reduced to about 40% (Fig. 1B; vehicle: 100 ± 2.64 %; 6-OHDA: 36.85 ± 3.46%; p < 0.0001, Mann-Whitney test). We also carried out the unbiased stereology for TH-positive SN neurons in the late phase and detected a very similar degree of loss (Suppl. Fig. 1C, vehicle: 102.6 ± 7.9 %; 6-OHDA: 39.96 ± 3.7 %; p = 0.0262, Mann-Whitney test; please add N numbers). Importantly, these results indicated that SN DA cell loss was non-progressive throughout our observation period of more than 2 months. In addition, we analyzed the axonal compartment of DA midbrain neurons by determining striatal TH-optical densities, both for early and late phase (Suppl. Fig. 1). Similar to the midbrain DA cell body counts, we found a stable reduction of about 50% of TH-immunosignal in the ipsilateral DS, again normalized to the contralateral side (Suppl. Fig. 1A 21^st^ day – vehicle: 98.4 ± 2.2 %; 6-OHDA: 42.5 ± 3.0 %; Suppl. Fig. 1B > 64 days – vehicle: 94.4 ± 2.7 %; 6-OHDA: 48.3 ± 3 %), both in early and late phase. By comparison, the ventral striatum was only mildly affected (ca. 20 % reduction, Suppl. Fig. 1A 21^st^ day – vehicle: 94.6 ± 2.0 %; 6-OHDA: 81.4 ± 3.0 %; Suppl. Fig. 1B > 64 days – vehicle: 97.8 ± 2.0 %; 6-OHDA: 75.2 ± 5.0 %). In summary and in accordance with previous studies, our model induced stable 60% loss of SN DA neurons, associated with a stable 50% reduction of DS TH-immunoreactivity throughout the entire observation period (Bez, Francardo, and Cenci 2016; Cenci and Björklund 2020; Schwarting and Huston 1996). This stability provided a suitable framework to identify time-dependent homeostatic changes in surviving SN DA neurons.

Fig 1 C shows the data from continuous behavioral monitoring of unilateral 6-OHDA and vehicle-treated mice before and up to 68 days post-infusion. Based on previous studies (Cenci and Björklund 2020; Schwarting and Huston 1996), we focused on the dynamics of drug-free spontaneous turning behavior during open field locomotion (Fig. 1C, lefts panels). In contrast to vehicle-infused mice, which displayed a stable symmetric ratio (ca. 50%) of ipsi-to contral-lateral turning throughout the entire experiment, 6-OHDA-infused mice showed a dramatic shift to ipsilateral turning immediately after treatment (Figure 1C, right panel). Interestingly, ipsilateral turns occurred in long sequences of up to 40 individual turns, a pattern not observed in controls (Suppl. Fig. 3). Importantly, these turning sequences as well as the overall ipsilateral bias gradually recovered completely over a 2-months post-lesion period (two-way-ANOVA, p-value across time p<0.0001, p-value across groups p<0.0001, significant difference between vehicle and 6-OHDA group till day 40, Šídák’s multiple comparisons test). Analysis of turning features, such as diameter or velocity, revealed that the recovered contralateral turns were similar to turns performed by vehicle-treated mice (Suppl. Fig. 4). Also, open field locomotion, quantified as total track length per session, recovered almost completely (Suppl. Fig. 1D). However, other motor behavior, the contralateral paw use during spontaneous rearing in the cylinder test, did not show significant recovery throughout the experimental period (Fig. 1D) (21st post-infusion day: vehicle 38.5 ± 5.2%, 6-OHDA 5.7 ± 1.4%, p < 0.0001, Mann-Whitney test; 68th post-infusion day: vehicle 63.7± 6.6%, 6-OHDA 3.2 ± 1.8%, p < 0.0001, Mann-Whitney test). Finally, repeating the continuous behavioral monitoring and the quantification of TH immunohistochemistry in midbrain and striatum with female C57Bl6N mice, infused with either 6-OHDA or vehicle, showed very similar results and gave no indications for sex differences (Suppl. Fig. 2).

To investigate the functional properties in immunohistochemically-identified surviving SN DA neurons, we performed *in vivo* extracellular recordings combined with juxtacellular neurobiotin (NB) labeling and *in vitro* whole-cell patch-clamp recordings combined with neurobiotin filling for two timepoints: at an early timepoint (21st day after lesion, early phase), where a strong turning asymmetry was present, and at a later time period (> 64 days post-lesion, late phase) when turning symmetry had recovered.

### Surviving SN DA neurons in the early post-6-OHDA phase exhibit impaired firing properties both *in vivo* and *in vitro*

At the early post-6-OHDA phase, we explored the *in vivo* spontaneous activity of surviving identified SN DA neurons in isoflurane-anesthetized mice, using single-unit extracellular recordings combined with juxtacellular neurobiotin labelling and post-hoc TH immunohistochemistry (see schema Fig. 2A). Figure 2B shows the *in vivo* extracellular electrical activity (top panel) and identification of a representative SN DA neuron (lower right panel) from a control mouse, 3 weeks after vehicle infusion. Note the typical combination of irregular single spike firing with the occurrence of transient fast burst discharges (bursts were defined by the 80/160ms Grace criteria; see methods for details) and longer pauses. The interspike interval (ISI) histogram, which captures ISIs from the entire recording time of 10 min for this cell, displays the typical features of an *in vivo* bursty DA neuron, i.e. a bimodal ISI distribution representing intra– and interburst intervals over a broad dynamic range between ∼30-1000 ms (lower left panel). This recorded and labelled cell was localized in the medial SN and characterized as TH-immunopositive. In comparison, Fig. 2C displays the representative *in vivo* activity of a surviving and identified SN DA neuron 3 weeks after 6-OHDA-infusion. Here, while irregular single-spike pacemaking was present similar to controls, fast burst events and slow pauses were almost completely absent (top panel). Indeed, the ISI histogram of this neuron, for the entire 10 min of recording, showed a compressed dynamic firing range and was well-described by a unimodal, Gaussian-like distribution without fast intraburst and slow interburst intervals (Fig. 2C, lower left panel). This recorded and labelled cell was also localized in the medial SN and identified as TH-immunopositive.

**Figure 2.**
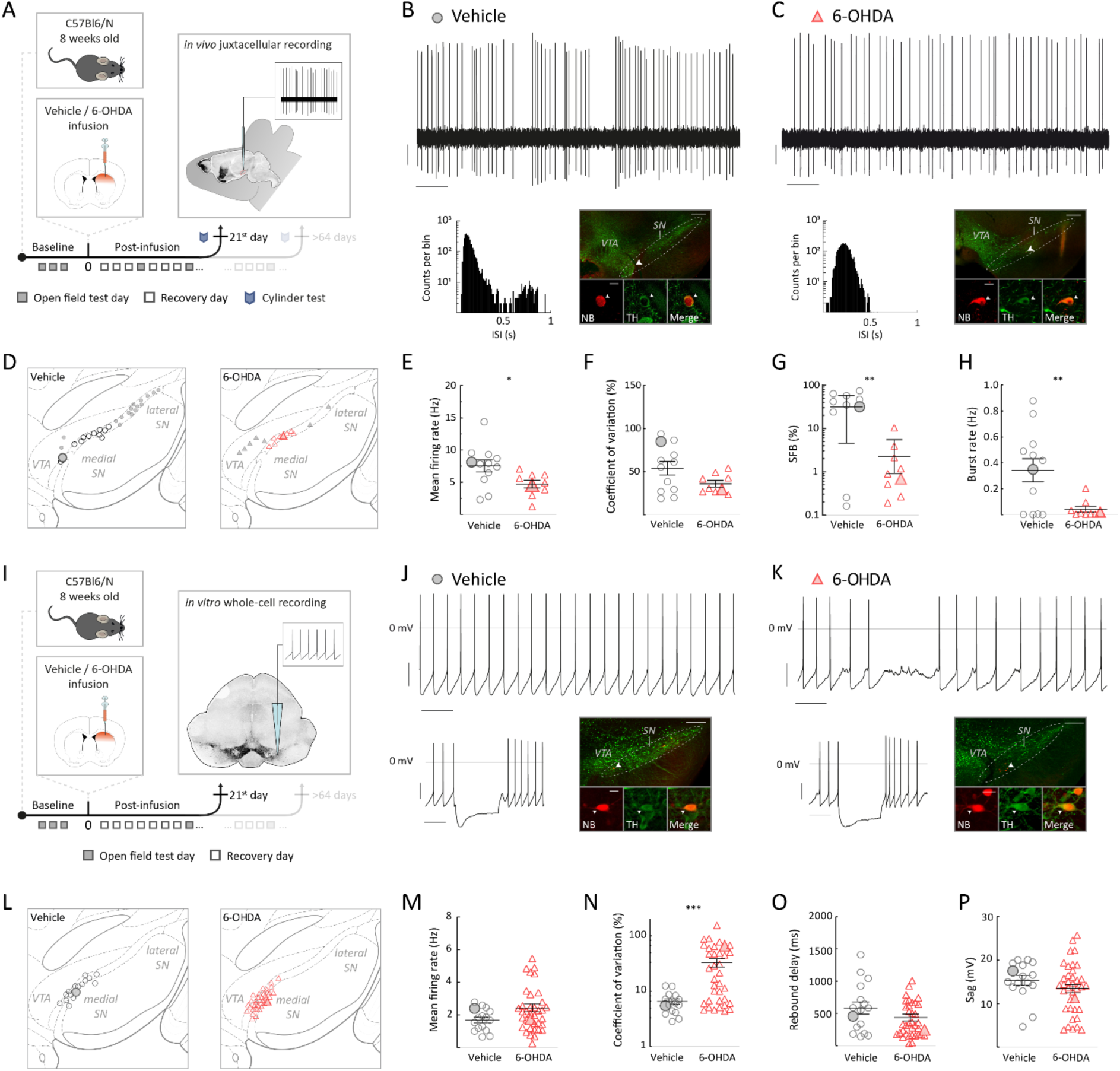
Surviving SN DA neurons at early post-6-OHDA phase exhibited a compressed dynamic range with a 10-fold decrease in *in vivo* bursting and a 5-fold decree in *in vitro* pacemaker regularity (A), (I) Experimental design, illustrating timeline of behavioral assays, followed by terminal *in vivo* juxtacellular recordings (A) or by terminal *in vitro* whole-cell recordings (I) at the 21^st^ post-infusion day. (B), (C) Top: 10s original recording trace of spontaneous *in vivo* single-unit activity from SN DA neurons in vehicle (B) and 6-OHDA-infused mouse (C). Scalebars: 1s, 0.2mV. Below, left: corresponding ISI histograms. Below, right: corresponding confocal images of juxtacellularly labelled and immunohistochemically identified DA neuron. Note the sparse bursting of the surviving SN DA neuron from the 6-OHDA-infused mouse at early phase. (D), (L) Anatomical mapping of all extracellularly recorded and juxtacellularly labelled DA neurons (D), and of all in vitro recorded and filled DA neurons (L), projected to bregma –3.08 mm. Location of example SN DA neurons in (B), (C), (J), (K) are highlighted. Smaller symbols represent DA neurons that have been recorded and identified, but not included in the group data analysis in (E-H). (E-H) Scatter dot-plots, showing significant decrease of *in vivo* mean firing rate (E), percentage of SFB (G) and burst rate (H) and no significant differences in CV (F) between the vehicle and 6-OHDA-infused mice. Note the 10-fold decrease in SFB for 6-OHDA-infused mice. (J), (K) Top: 10s original recording trace of *in vitro* whole-cell recording of spontaneous SN DA neuron activity in a vehicle (J) and 6-OHDA-infused (K) mouse. Scalebars: 1s, 20mV. Note that the 6-OHDA DA neuron has a highly irregular pacemaking. Below, left: corresponding to hyperpolarizing current injection. Below, right: confocal images of NB filled and immunohistochemically identified DA neuron. (M-P) Scatter dot-plots, showing no difference in *in vitro* mean firing rate (M), rebound delay (O) or sag-component (P) and a 5-fold increase in CV (N). Immunohistochemical imaging for all four DA neurons are displayed in 10x and 60x magnifications (green, TH; red, NB), scalebars: 200µm, 20µm. All data are presented as mean ± standard error of mean (SEM).

All *in vivo* recorded DA neurons, like the two representative cells shown above, from the early phase were successfully juxtacellularly labeled with NB, post-hoc immunohistochemically identified and localized within the ventral midbrain (Fig 2D, vehicle: n =28, N =15; 6-OHDA: n =18, N =16). For best comparison, we restricted our analysis to those DA midbrain neurons localized in the medial half of the SN (mSN), where it was possible to sample a sufficient number of surviving and spontaneously active DA neurons after 6-OHDA lesion. In contrast, we only managed to detect a small number of active DA neurons in the lateral SN (lSN), the most vulnerable part of the DA midbrain (Figure 2D). Consequently, Figure 2 E-H compares the group data of surviving mSN DA neurons at early phase between vehicle and 6-OHDA treatment. We detected an about 60% reduction of the mean firing rates for post-6-OHDA mSN DA neurons compared to vehicle-treated controls (Fig.2E; vehicle: mean frequency = 7.5 ± 0.9 Hz, 6-OHDA: mean frequency = 4.7 ± 0.6 Hz, p = 0.0184, Mann-Whitney test) and no significant differences in overall firing regularity (Fig. 2F; vehicle: CV = 53.9 ± 7.8 %, 6-OHDA: CV = 36.1 ± 3.7 %, p = 0.1694, Mann-Whitney test). Moreover, we observed a significant 10-fold reduction of bursting – expressed as the percentage of spikes fired in bursts (SFB) and burst rate – in mSN DA neurons in the 6-OHDA group compared to vehicle controls (Fig. 2G, vehicle: SFB = 31.2 ± 7.7; 6-OHDA: SFB = 2.2 ± 1.1 %; p = 0.0032, Welch’s t test; Fig. 2H, vehicle: burst rate = 0.34 ± 0.09 Hz, 6-OHDA: burst rate = 0.04 ± 0.02 Hz; p = 0.0066, Welch’s t test).

All other analyzed firing parameters, such as intra-burst firing frequencies, identified no additional differences between the two groups (Suppl. Fig. 5A-G). In summary, we detected a rightward shift to lower firing frequencies with a corresponding compression of the dynamic *in vivo* firing range in mSN DA neurons surviving the 6-OHDA lesion (Suppl. Fig. 5I). Thus, in addition to DA cell loss, the dysfunction of surviving DA neurons might be a novel contributing factor to the extensive motor impairment, observed during the early post-6-OHDA phase. In particular, transient bursts of DA SN neurons have been shown to provide start and stop signals for motor control (see discussion for details).

Burst discharges in DA neurons are orchestrated by the interplay of patterned synaptic inputs with their intrinsic excitability, giving rise to different types of *in vivo* bursting (Otomo et al. 2020). To identify a potential contribution of cell-autonomous changes in intrinsic excitability of surviving mSN DA neurons to their impaired *in vivo* dynamics, we also studied these cells *in vitro* in synaptic isolation (see Schema 2I). Figure 2J shows a representative, stable and regular pacemaker activity of an identified mSN DA neuron from a vehicle-infused control (lower right panel), as well as a subthreshold response to hyperpolarizing current injection (lower left panel). In contrast, pacemaker activity of mSN DA neurons post-6-OHDA infusion was unstable and characterized by intermittent periods of action potential failure (see Fig. 2K, upper panel). However, subthreshold responses were similar to controls (lower right panel). Analogous to the *in vivo* experiments, all *in vitro* recorded DA neurons were NB filled, identified, and mapped, resulting in a similar medial localization of surviving SN DA neurons (Fig. 2L). When *in vitro* pacemaker properties were compared between identified mSN DA neurons from the vehicle and the 6-OHDA groups (Fig. 2L), we detected no differences in mean firing rates (Fig. 2M, vehicle: firing rate = 1.7 ± 0.2 Hz, 6-OHDA: firing rate = 2.4 ± 0.3 Hz, p = 0.1448, Mann-Whitney test), rebound delays or sag-amplitudes (Fig. 2O, vehicle: rebound delay = 584.9 ± 95.8 ms, 6-OHDA: rebound delay = 429.6 ± 56.1 ms, p = 0.1738, Mann-Whitney test; Fig. 2P, vehicle: sag-amplitude = 15.3 ± 1.1 mV, 6-OHDA: sag-amplitude = 13.5 ± 0.96 mV, p = 0.1395, Mann-Whitney test). In contrast, a significant, about 5-fold increase of pacemaker irregularity expressed as CV was detected (Fig.2N, vehicle: CV = 6.5 ± 0.7 %; 6-OHDA: CV = 33.9 ± 5.8; p = 0.0007, Mann-Whitney test). Detailed observation of the raw traces revealed that the increased CV was mainly mediated by a combination of intermittent phases of firing failure, similar to the one shown in Fig. 2K, and episodes of higher frequency firing (as quantified in the ISI distribution histogram, Suppl. Fig. 6H). We also showed that these differences were preserved both in on-cell and whole-cell recordings indicating the latter did not induce a differential bias in any of the two groups (e.g. different calcium buffering) (Suppl. Fig. 6F-G).

In short, this data demonstrates functional alterations in DA SN neurons surviving three weeks after a 6-OHDA infusion. These neurons exhibited a compressed dynamic range, including a 10-fold reduction of *in vivo* bursting. Furthermore, we observed a 5-fold decrease of *in vitro* cell-autonomous pacemaker stability with intermittent failure of action potential firing. This compromised intrinsic *in vitro* excitability likely contributes to the 10-fold reduction of high frequency *in vivo* firing.

In order to address the question of whether these altered properties represent stable adaptations or reflect a transient state of damage, we also studied *in vivo* and *in vitro* electrophysiology of surviving mSN DA neurons in the late phase after lesion, characterized by partial behavioral recovery of motor functions.

### Surviving mSN DA neurons in the late post-6-OHDA phase display full recovery of the *in vivo* dynamic range associated with an accelerated *in vitro* pacemaker

We recorded and analyzed surviving mSN DA neurons in the late 6-OHDA phase analogous to the early phase (compare Fig. 3A/I and Fig. 2A/I). In contrast to the early phase, we found no significant differences in the electrophysiological *in vivo* properties between the post-lesional and vehicle-infused groups in the late phase (compare Fig 3B with Fig. 3C). Among others (Suppl. Fig. 7), this implies that mean firing frequencies, dynamic ranges, CV, SFB and burst rates did not differ in chronically surviving mSN DA neurons compared to those in controls (Fig. 3E, vehicle: mean firing rate = 6.9 ± 0.8 Hz; 6-OHDA: mean firing rate = 7.5 ± 1.1 Hz; p = 0.6038, Mann-Whitney test; Fig. 3F, vehicle: CV = 54.9 ± 12.2%; 6-OHDA: CV = 53.3 ± 12.6%, p > 0.9999, Mann-Whitney test; Fig. 3G, vehicle: SFB = 20.2 ± 9.1%, 6-OHDA: SFB = 22.2 ± 8.9%, p = 0.8784, Mann-Whitney test; Fig. 3H, vehicle: burst rate = 0.17 ± 0.08 Hz; 6-OHDA: burst rate = 0.30 ± 0.15 Hz, p = 0.4308, Mann-Whitney test). This suggests that surviving mSN DA neurons show full recovery of their electrophysiological properties within 2-months after the lesion. However, our *in vivo* results alone cannot distinguish between two scenarios: first, a slow functional recovery process that eventually leads to return of the physiological activity range or second, the return to the full dynamic firing range *in vivo* is more than simple repair to the *status quo ante* but involves a homeostatic adaptation of the functional properties of the DA neuron itself. To resolve this question, we also recorded the *in vitro* pacemaker properties of the surviving DA mSN neurons in the late post-6-OHDA phase. Here, we found clear evidence for allostatic adaptation of these DA neurons (i.e the presence of new homeostatic setpoint). When comparing intrinsic pacemaker frequencies, we noticed that late 6-OHDA survivors discharged almost 2-fold faster compared to vehicle-controls (compare Fig. 3J with 3K, Fig. 3M vehicle: firing rate = 1.7 ± 0.2 Hz; 6-OHDA: firing rate = 2.7 ± 0.2 Hz, p = 0.0035, Mann-Whitney test). In contrast to speed, no significant difference in pacemaker regularity was observed (Fig. 3N, vehicle: CV = 9.19 ± 1.30 %, 6-OHDA: CV = 16.10 ± 6.66 %, p = 0.1245, Mann-Whitney test). Also, the sag components and the rebound delay were not significantly different (Fig 3O, vehicle: rebound delay = 512.8 ± 93.8 ms, 6-OHDA: rebound delay = 375.0 ± 58.9 ms, p = 0.1994, Mann-Whitney test; Fig. 3P, vehicle: sag-amplitude = 16.4 ± 1.2 mV, 6-OHDA: sag-amplitude = 19.1 ± 0.9 mV, p = 0.0528, Mann-Whitney test). Other *in vitro* spike features were also not significantly different between the two groups (Suppl. Fig. 8). This accelerated pacemaker phenotype was also observed in metabolically intact DA neurons recorded in the on-cell configuration but further accelerated in the whole-cell mode (Suppl. Fig. 8F-G).

**Figure 3.**
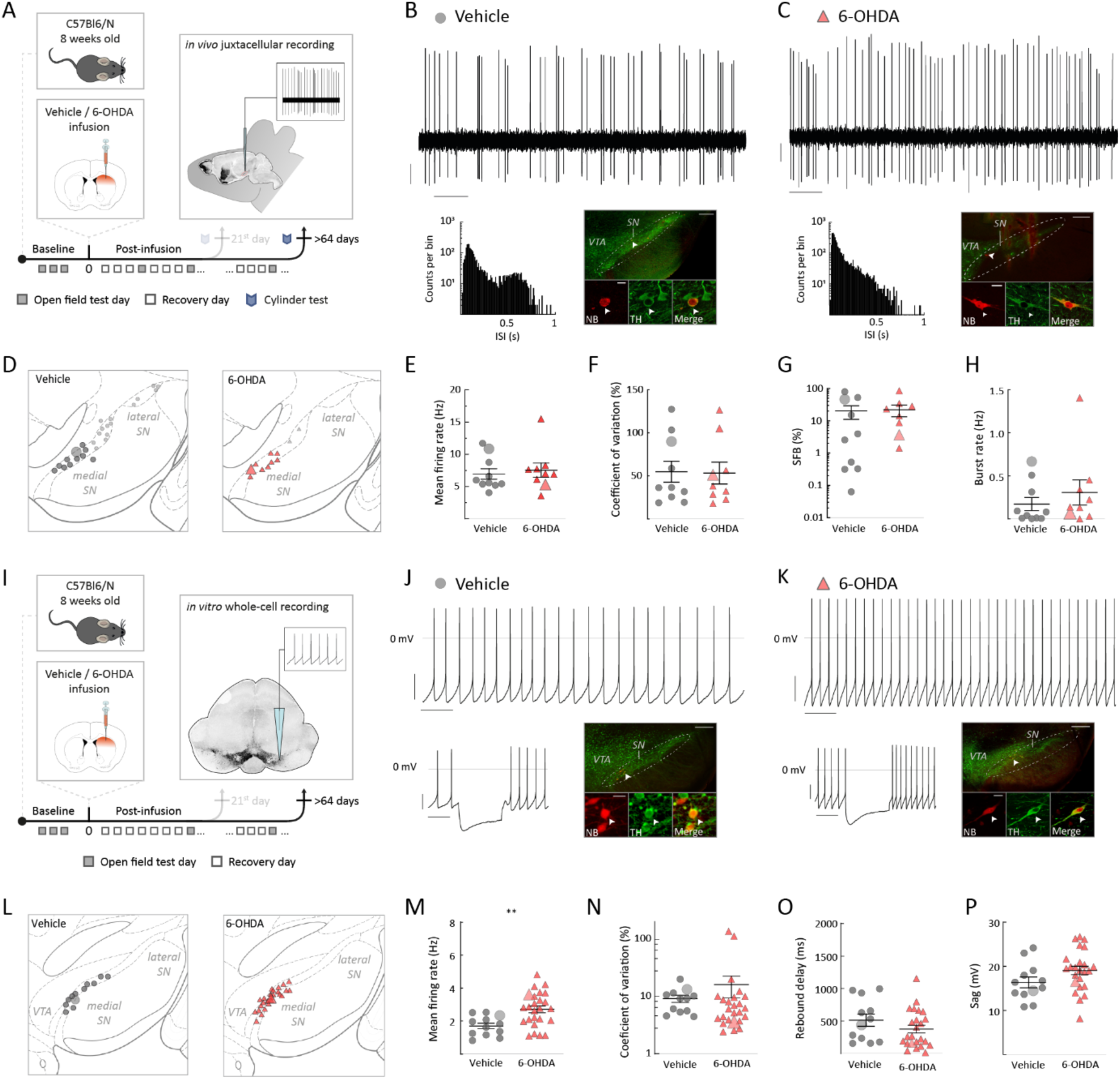
Surviving SN DA neurons at late 6-OHDA phase recovered *in vivo* burst firing and doubled their intrinsic pacemaker frequency *in vitro*. (A), (I) Experimental design, illustrating timeline of behavioral assays, followed by terminal *in vivo* juxtacellular recordings (A) or by terminal *in vitro* whole-cell recordings (I) after >64 post-infusion days. (B), (C) Top: 10s original recording trace of spontaneous *in vivo* single-unit activity from SN DA neurons in vehicle (B) and 6-OHDA-infused mouse (C). Scalebars: 1s, 0.2mV. Below, left: corresponding ISI histograms. Below, right: corresponding confocal images of juxtacellularly labelled and immunohistochemically identified DA neuron. (D), (L) Anatomical mapping of all extracellularly recorded and juxtacellularly labelled DA neurons (D), and of all *in vitro* recorded and filled DA neurons (L), projected to bregma –3.40 mm (D) and –3.08mm (L). Location of example SN DA neurons in (B), (C), (J), (K) are highlighted. Smaller symbols represent DA neurons that have been recorded and identified, but not included in the group data analysis in (E-H). (E-H) Scatter dot-plots, showing no significant difference in mean firing rate (E), CV (F), SFB (G) and burst rate (H). (J), (K) Top: 10s original recording trace of *in vitro* whole-cell recording of spontaneous activity from SN DA neurons in a vehicle (J) and 6-OHDA-infused (K) mouse. Scalebars: 1s, 20mV. Note that the 6-OHDA DA neuron has enhanced, but regular pacemaking. Below, left, corresponding to hyperpolarizing current injection. Below, right, confocal images of NB filled and immunohistochemically identified DA neuron. (M-P) Scatter dot-plots, showing doubling of the firing rate (M), no difference in CV (N), decrease in rebound delay (O) and increase of sag-component (P) for the 6-OHDA-treated mice in comparison to vehicle group. Immunohistochemical imaging for all four DA neurons are displayed in 10x and 60x magnifications (green, TH; red, NB), scalebars: 200µm, 20µm. All data are presented as mean ± standard error of mean (SEM).

In summary, functional recovery of the *in vivo* dynamic firing range of surviving mSN DA neurons was associated with homeostatic plasticity of pacemaking (i.e. an allostatic shift of pacemaker setpoint). Surviving mSN DA neurons not only recovered from instable pacemaking at the early phase but doubled their discharge rates in late phase.

### Accelerated pacemaker caused by downregulation of Kv4.3 channels in late surviving mSN DA neurons

As Kv4.3 A-type channels have been shown to be powerful regulators of DA pacemaker frequency (Subramaniam, Althof, et al. 2014; Subramaniam, Kern, et al. 2014; Liss et al. 2001; Khaliq and Bean 2008; Tarfa, Evans, and Khaliq 2017), we tested their potential role for the observed pacemaker acceleration in late phase DA mSN neurons by pharmacological occlusion experiments using 1µM AmmTx3, a selective Kv4.3 channel blocker (see scheme Fig. 4A). The presence of 1µM AmmTx3 not only accelerated discharge rates in both identified mSN DA neurons from vehicle– and 6-OHDA-infused mice (see Fig. 4BC for representative cells; Fig. 4D for mapping) but, importantly, also eliminated significant rate differences between the two treatment groups (Fig. 4E, vehicle: mean firing rate = 4.5 ± 0.2 Hz, 6-OHDA: mean firing rate = 4.8 ± 0.3 Hz, p = 0.285; Fig. 4 F, vehicle: CV = 7.8 ± 1.3 %, 6-OHDA: CV = 5.5 ± 0.4 %, p = 0.2517; both Mann-Whitney test). This result demonstrated that differences in Kv4.3 channel function were the main driver of the late post-6OHDA accelerated pacemaker phenotype. In addition, 1µM AmmTx3 also abolished the more subtle differences in rebound delays and sag amplitudes (Fig. 4G, vehicle: rebound delay = 31.7 ± 2.4 ms, 6-OHDA: rebound delay = 28.4 ± 4.0 ms, p = 0.0952; Fig. 4H. vehicle: sag-amplitude = 15.6 ± 1.1 mV, 6-OHDA: sag-amplitude = 16.4 ± 0.9 mV, p = 0.7434; both Mann-Whitney test, Suppl. Fig. 9). In summary, these experiments demonstrated that reduced Kv4.3 channel function in post-6-OHDA late phase DA mSN neurons had a causal role in the phenotype of accelerated *in vitro* pacing.

**Figure 4.**
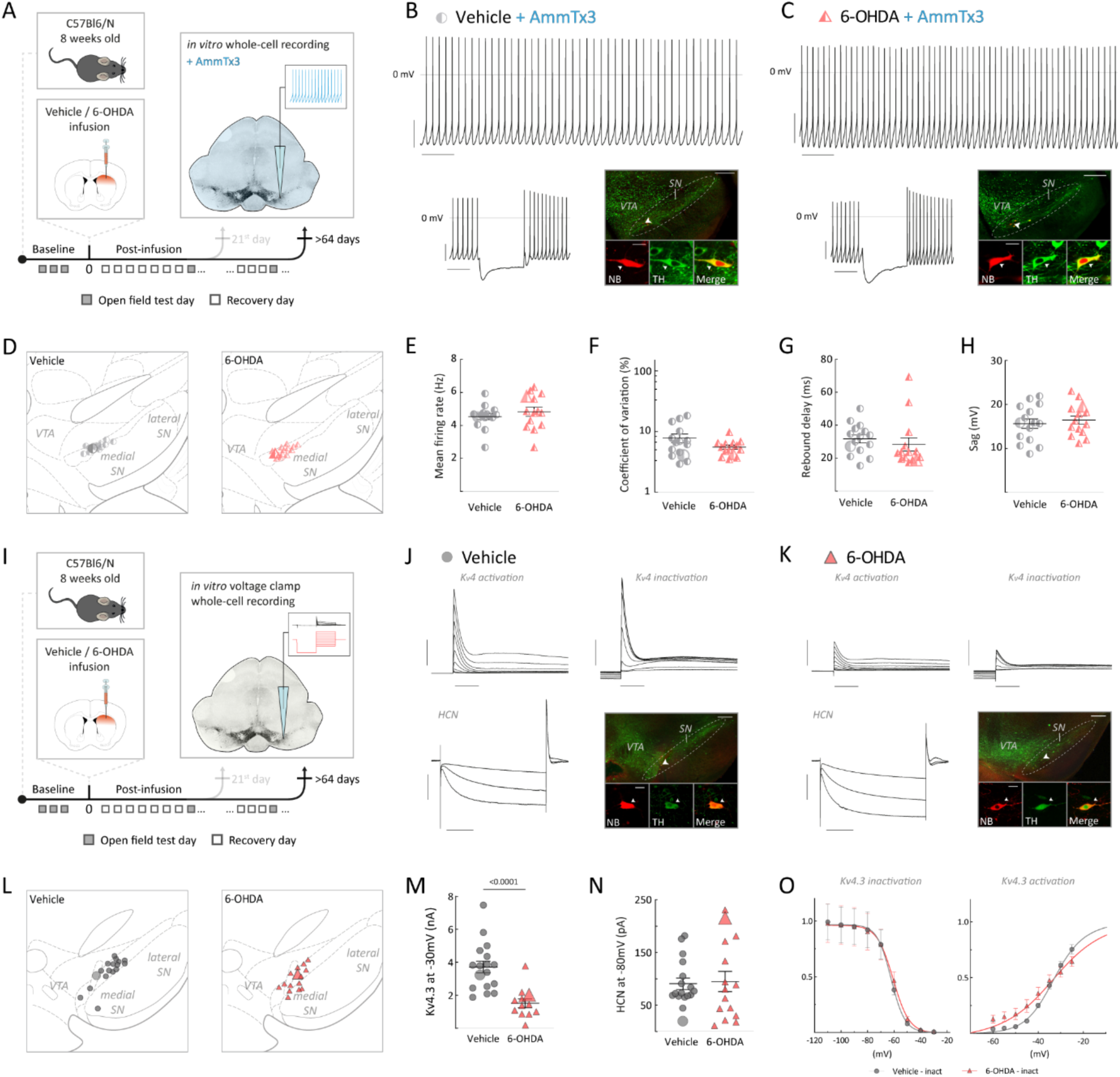
Enhanced pacemaker frequency of SN DA neurons at the late post-6-OHDA phase was mediated by K_v_4.3 channel downregulation. (A), (I) Experimental design, illustrating timeline of behavioral assays, followed by terminal *in vitro* whole-cell recordings under K_v_4.3 channel blocker AmmTx3 (A) or *in vitro* voltage-clamp whole-cell recordings (I) after >64 post-infusion days. (B), (C) Top: 10s original recording traces of *in vitro* whole-cell recording of spontaneous activity from DA SN neurons in AmmTx3-preincubated slices for a vehicle (B) and 6-OHDA-infused (C) mouse. Below, left: corresponding hyperpolarizing current injection. Scalebars: 1s, 20mV. Below, right: confocal images of NB filled and immunohistochemically identified DA neuron. (D), (L) Anatomical mapping of all in vitro recorded and filled DA neurons, projected to bregma –2.92 mm (D) and –3.16mm (L). (E-H) Scatter dot-plots, showing no differences in mean firing rate (E), coefficient of variation (F), rebound delay duration (G) and sag amplitude (H). (J), (K) Original recording trace of in vitro voltage-clamp recordings from DA SN neurons in a vehicle (J) and 6-OHDA-infused (K) mouse. Top, right: Zoom in from a K_v_4.3 channel activation. Top, left: Zoom in from a K_v_4.3 channel inactivation. Below, right: Zoom in from HCN-channel activation. Below, right, confocal images of NB filled and immunohistochemically identified DA neuron. Scalebars first row: 3nA, 100ms. Scalebars below, left: 5nA, 500ms. Note the small K_v_4.3 channel in/activation peak in a surviving DA neuron from a 6-OHDA mouse in comparison the one from an ACSF treated mouse. Immunohistochemical imaging for all four DA neurons are displayed in 10x and 60x magnifications (green, TH; red, NB), scalebars: 200µm, 20µm. All data are presented as mean ± standard error of mean (SEM). (M) Scatter dot-plots, showing a significant, half of maximum K_v_4.3 channel conductance in surviving DA neurons. (N) Scatter dot-plots, showing no difference in HCN channel delta current (I_peak_ – I_steady_) between groups. (O) Normalized K_v_4.3 current (I_peak_ – I_steady_) at different voltage steps, resulting in inactivation and activation curve for both groups. All data are presented as mean ± standard error of mean (SEM).

To directly quantify the gating properties of transient outward (A-type) potassium currents mediated by Kv4.3 channels, we carried out whole-cell voltage-clamp recordings (see scheme Fig. 4I-O). As reported before (Liss et al. 2001), voltage-steps from negative holding potentials to the relevant subthreshold range (e.g. –60 mV to –30 mV) activated large, fast inactivating outward currents in the range of 1-10 nA in vehicle injected controls. In contrast, the same voltage steps activated smaller fast-inactivating outward currents in post-6-OHDA late phase DA mSN neurons. These experiments demonstrated about 50% smaller maximal A-type Kv4.3 potassium currents in late surviving DA mSN neurons from 6-OHDA-treated mice compared to vehicle-treated controls (Fig. 4 M. vehicle: Kv4.3 channel current at –30mV = 3.8 ± 0.4 nA, 6-OHDA: Kv4.3 channel current at –30mV = 1.7 ± 0.3 nA, p < 0.0001, Mann-Whitney test). When comparing voltage-dependent gating parameters, we found no changes in the biophysical channel properties apart from differences in the slope of the steady-state activation curves (Fig. 4M, Suppl. Fig. 10A-C).

By comparison, recordings of HCN currents in whole-cell voltage-clamp recordings showed no differences in current amplitudes between the groups (Fig. 4N, vehicle: ΔHCN = 90.8 ± 10.7 pA, 6-OHDA: ΔHCN = 94.9 ± 19.1 pA, p = 0.7375, Mann-Whitney test). These results substantiated the selective role of reduced Kv4.3 function in causing the accelerated pacemaker in the post-6-OHDA late phase surviving mSN DA neurons. As the reduction of Kv4.3 function might be caused by a number of different mechanisms, including e.g. reduced protein expression and/or membrane delivery of Kv4.3 subunits as well as post-translational modifications of existing Kv4.3 channel complexes (e.g. phosphorylation, redox-modification), we decided to assess Kv4.3 protein expression via immunohistochemistry.

We performed Kv4.3 immunohistochemistry and confocal imaging in vehicle and 6-OHDA treated mice both on the ipsilateral and contralateral infusion side (see scheme in Fig. 5A). We defined DA-selective ROIs by using TH-immunopositive areas to determine the intensity of Kv4.3-immunosignals exclusively within DA neurons (i.e. TH-signal based mask) (Subramaniam, Althof, et al. 2014). Figure 5B and 5C compare Kv4.3-immunoreactivities in the midbrain between the contralateral control side and the affected ipsilateral side during the late post-6-OHDA phase. Ipsilateral post-lesional TH-positive neurons showed reduced Kv4.3-immunosignals compared to the control side. Semi-quantitative comparison of TH-ROIs of the entire SN between contralateral control side and 6-OHDA-treated side revealed a significant reduction of average Kv4.3-immunosignal on the 6-OHDA treated side of about 25% (Figure 5D, contralateral side n = 4004, ipsilateral side, n = 2645, D = 0.2014, p < 0.0001, two-sample Kolmogorov–Smirnov test). A more detailed analysis of regional differences within the SN revealed that the reduction in Kv4.3 immunoreactivity was significant for the medial SN (Figure 5E, contralateral side, n = 4004: 73.03 ± 0.39, ipsilateral side, n = 2645: 54.8 ± 0.49, p < 0.0001, unpaired t-test), where we recorded Kv4-currents, but even more pronounced in the lateral parts of the SN (Suppl. Fig. 11C). As Kv4.3 channel subunits are expected to be mainly located on the cell membrane, we also investigated whether Kv4.3-immunosignals were differentially distributed across cell compartments. A robust, about 17% reduction of Kv4.3-immunosignals was observed in ipsilateral post-lesional TH-positive neurons both for cell membrane and cytoplasm ROIs (Suppl. Fig. 11D). In contrast, analogous quantification in vehicle-treated mice revealed no contra-ipsilateral differences in Kv4.3-immunoreactivity (Suppl. Fig. 11E-F). In summary, these immunohistochemical experiments provided evidence for a reduced expression of Kv4.3 channel proteins in late post-6OHDA mSN DA neurons. Reduced Kv4.3 protein expression is consistent with a reduction of functional Kv4.3 channels in surviving mSN DA neurons as described above and suggests that pacemaker acceleration is mediated by a reduced number of Kv4.3 channels.

**Figure 5.**
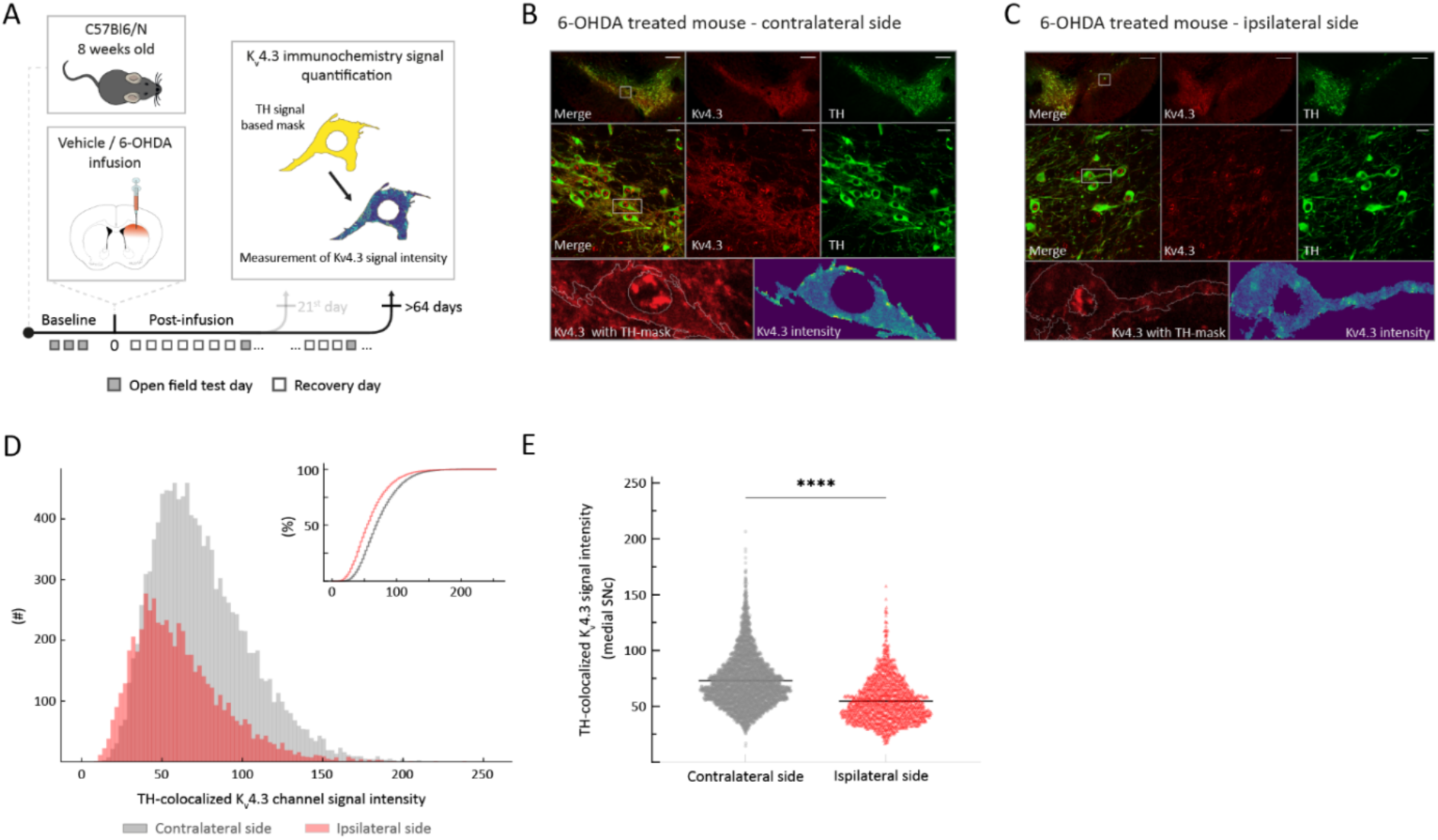
Surviving DA SN neurons at the late post-6-OHDA phase showed lower. **K**_v_**4.3 channel immunohistochemical signal** (A) Experimental design, illustrating timeline of behavioral assays, followed by multi-labeling immunohistochemistry for exploring K_v_4.3-channel expression differences after >64 post-infusion days. (B), (C) Top: 4x magnification of midbrain of a 6-OHDA-infused mouse, >64 days post-lesion – contralateral side (B), and corresponding ipsilateral side (C). Middle: 60x magnification in the highlighted area from 4x image (green, TH; red, K_v_4.3). Bottom, left: zoom-in on an example ROI (highlighted in 60x image). Bottom, right: color-coded Kv4.3-channel immunohistochemical signal intensity in the example ROI. Note K_v_4.3-channel signal decrease in surviving DA neurons on the ipsilateral to the injection side. (D) Histogram showing intensity of K_v_4.3 immunosignals for all TH-positive ROIs, from ipsilateral, lesioned, side (in red) and from contralateral side (in grey). Inset, same data shown as a cumulative distribution. Note a clear right-shift to lower intensities for the ipsilateral side. (E) Comparison of mean TH-colocalized K_v_4.3 immunosignals from medial SN from ipsilateral, lesioned, side, and contralateral, as control side. All data are presented as mean ± standard error of mean (SEM).

## Discussion

Our study provides the first combined *in vitro* & *in vivo* electrophysiological characterization of identified DA neurons in the substantia nigra surviving a partial 6-OHDA lesion. Studying these surviving SN DA neurons at two time points, we discovered time-dependent post-6-OHDA-selective differences of firing properties both *in vivo* and *in vitro*. Early after lesion, and coinciding with prominent behavioral impairments, we detected a selective and dramatic (about 10-fold) reduction of *in vivo* burst firing in identified surviving SN DA neurons. This is reminiscent of an early study by Hollermann & Grace (Hollerman and Grace 1990), who reported reduced bursting in putative SN DA neurons one week after a partial lesion. Considering the Berretta et al 2005 study (Berretta et al. 2005), these early electrophysiological effects may reside from 6-OHDA triggering a cascade of events, e.g. hyperpolarization via activation of K-ATP channels. At this early time point, we also found unstable and more irregular *in vitro* pacemaker activity in these cells, with no differences in mean firing rates, compared to those from vehicle-infused controls.

Conversely, at a later post-lesion time point (>2 months), when partial motor recovery had occurred, we detected no differences in the *in vivo* firing range – including high frequency bursts– between surviving SN DA neurons post-6-OHDA and those from vehicle-infused mice. Surprisingly, the *in vitro* pacemaker rates in post-6-OHDA DA survivors were not only stable but almost twice as fast compared to those from vehicle-controls. Finally, we pinpointed functional and protein Kv4.3 channel downregulation as the leading cause for this chronic post-lesional pacemaker acceleration, while the HCN channel function remained relatively stable. In essence, we identified a slow recovery of the dynamic *in vivo* firing range of surviving post-lesional DA SN neurons, linked to an allostatic acceleration of the intrinsic pacemaker by Kv4.3 channel downregulation.

Our results suggest a simple homeostatic scheme, where reducing the population size of DA SN neurons by half is at least in part compensated by a two-fold increase of pacemaker speed in the remaining half of the DA SN cell population. Given the 10-fold reduction of *in vivo* bursting in the initial period after the lesion, it further suggests that the pacemaker setpoint rate is coupled to the *in vivo* high frequency burst rate in DA mSN neurons. It is possible that a burst-related calcium signal might be detected and integrated over time to control pacemaker rate via e.g. tuning Kv4.3 expression. Recent studies have implicated the mitochondrial calcium uniporter (MCU) in coupling bursts to mitochondrial Ca^2+^, acting as a calcium sensor for neuronal firing rate homeostasis (Katsenelson et al. 2022). Future mechanistic studies will be necessary to investigate how *in vivo* bursting and intrinsic pacemaking are coupled in distinct DA subpopulations. It is however interesting to note that the very first descriptive study of in vivo SN DA firing patterns had already found a positive correlation between the degree of in vivo burst firing and single spike firing rate (A. A. Grace and Bunney 1984). In agreement with this proposal, it was also previously found that an about 4-fold-reduction of *in vivo* burst rate via DA-selective NR1 knockout resulted in a scaled reduction of *in vivo* firing rate (Zweifel et al. 2009).

In contrast to the homeostatic role of flexible Kv4.3 expression identified in this study, we previously identified Kv4.3 channels in SN DA neurons as a pathophysiological target in a transgenic mutant (A53T-SNCA) α-synuclein mouse model (without cell loss) (Subramaniam, Althof, et al. 2014). In this model, we found a pacemaker acceleration caused by oxidative impairment of Kv4.3 channels. However, the activity of SN DA neurons in the α-synuclein model was accelerated also *in vivo.* Thus, mutant α-synuclein expression also caused a shift of the *in vivo* firing range of SN DA neurons towards higher frequency. While Kv4.3 subunits were downregulated at the protein level in post-lesional SN DA neurons, mutant α-synuclein induced oxidative Kv4.3 dysfunction as well as protein upregulation (Subramaniam, Althof, et al. 2014). It will be interesting to test in follow up studies if and how α-synuclein pathology affects firing and burst rate homeostasis in DA SN neurons.

Beyond pacemaker adaptations in surviving SN DA neurons, we assume that their chronic *in vivo* electrophysiological phenotype might also be reshaped by network level plasticity in response to dopamine-depletion.

Our finding that the mean *in vivo* discharge rates were not different to controls in the presence of an accelerated intrinsic pacemaker strongly suggests a shift of the synaptic excitation-inhibition (E-I) balance towards more inhibition. Numerous studies have found altered synaptic inhibition in the dopamine-depleted basal ganglia (recently reviewed in (Zhang et al. 2021)). In particular, Heo and colleagues recently demonstrated a chronic E-I balance shift toward more inhibition across several PD models including 6-OHDA (Heo et al. 2020). They identified a substantial contribution of additional GABA synthesis and release from reactive astrocytes in the midbrain. It has also been well-established that basal ganglia GABA neurons fire in more synchronized hence more effective fashion after DA depletion (Cagnan, Duff, and Brown 2015; Milosevic et al. 2018; Phillips et al. 2020; Tinkhauser et al. 2020; Wichmann, Bergman, and DeLong 2018; Evans et al. 2020). This has recently been confirmed by elegant *in vivo* single cell resolved studies using either gCAMP-based calcium monitoring or *in vivo* patch-clamp approaches (Ketzef et al. 2017; Kravitz et al. 2010; J. G. Parker et al. 2018; P. R. L. Parker, Lalive, and Kreitzer 2016; Sitzia et al. 2020). The accelerated pacemaker in response to enhanced net inhibition would also be expected within the framework of homeostatic plasticity (Turrigiano 2012).

Our study has several limitations. First, we explored a single-hit, non-progressive partial lesion model, which needs to be differentiated from *state-of-the-art* PD rodent models, which develop disease-relevant α-synuclein pathology and show progressive loss of DA SN neurons (Thakur et al. 2017). Second, we do not differentiate between DA SN subtypes. We recently showed that lateral SN DA neurons projecting to DLS possess a distinct *in vivo* firing phenotype compared to medial SN DA neurons projecting to either DLS or DMS in mice (Farassat et al. 2019). Therefore, recording from lateral DA SN population in lesion models would be important. However, in our current partial 6-OHDA lesion model, these lateral SN DA neurons were too severely lost to allow for more than anecdotal post-lesion electrophysiological analysis (see Fig. 3). A milder version of the current model would be needed to identify their properties and responses to partial lesion. Regarding the surviving DA neuron in the medial SN, we previously showed that they have distinct projection targets including DLS, DMS and lateral shell of nucleus accumbens (Farassat et al. 2019). In the current study, we have not aimed for identification of axonal projections to avoid potentially confounding effects of additional brain surgery. Nevertheless, this important issue should be addressed in follow-up studies by e.g. molecular subtyping approaches (Heymann et al. 2020; Poulin et al. 2018; Saunders et al. 2018). Third, we are aware that *in vivo* recordings carried out in isoflurane do not display the full dynamic spectrum of DA SN firing compared to awake, freely moving animals (we have discussed the aspect extensively in Farassat et al. 2019).

Finally, we would like to speculate about the possible implications of our findings for PD. In principle, cell-loss induced pacemaker plasticity, as identified here, might have a dual nature. On one hand, they might render the surviving DA SN neurons more robust in defending their phenotype and function. On the other hand, the allostatic pacemaker acceleration may amplify the innate vulnerability of an already at baseline metabolically and oxidatively challenged neuron type. The latter would be an extension of the “stressful pacemaker” hypothesis of DA vulnerability (Chan et al. 2007; Surmeier 2007; Surmeier, Obeso, and Halliday 2017; Surmeier, Halliday, and Simuni 2017) with its potential clinical implications(Guzman et al. 2018; Liss and Striessnig 2019; Ortner et al. 2017; Ortner 2021; The Parkinson Study Group STEADY-PD 2020). In summary, we identified the homeostatic capacity and pacemaker allostasis of surviving DA SN neurons in response to cell loss.

## Methods

### Animals

Male and female C57Bl6/N mice (Charles River Laboratories) were used for the study. The mice were 8 weeks old, group housed and maintained on a 12-hour light-dark cycle. All experiments and procedures involving mice were approved by the German Regierungspräsidium Darmstadt (V54-19c20/15-F40/30). In total, 152 mice were used for this study (see table below).

|  | Early phase |  | Late phase |  |
| --- | --- | --- | --- | --- |
|  | Mice (#) | Cells (#) | Mice (#) | Cells (#) |
| ***In vivo* juxta & behavior – vehicle** | 15 | 12 – medial/<br>28 – all | 12 | 10 – medial/<br>26 – all |
| ***In vivo* juxta & behavior – 6-OHDA** | 16 | 9 – medial/ 16<br>– all | 10 | 9 – medial/ 11<br>– all |
| ***In vitro* – vehicle** | 3 | 16 | 3 | 12 |
| ***In vitro* – 6-OHDA** | 6 | 37 | 5 | 25 |
| ***In vitro* & AmmTx3 – vehicle** | - | - | 3 | 15 |
| ***In vitro* & AmmTx3 – 6-OHDA** | - | - | 3 | 14 |
| ***In vitro* voltage clamp experiments – vehicle** | - | - | 3 | 17 |
| ***In vitro* voltage clamp experiments – 6-OHDA** | - | - | 3 | 15 |
| **Female behavior experiments – vehicle, 6-OHDA** | - | - | 10 | - |
| **Female histology – vehicle, 6-OHDA** | - | - | 8 | - |
| **6-OHDA dose response** | 12 | - | - | - |
|  | Mice (#) |  | Mice (#) |  |
| **Kv4.3 Immunocytochemistry** | 8 |  | 8 |  |
| **SN quantification – vehicle** | 9 |  | 3 |  |
| **SN quantification – 6-OHDA** | 9 |  | 3 |  |
| **Sum = 152** |  |  |  |  |

### Stereotactic 6-OHDA infusion

All surgeries were performed under general anesthesia in areflexic state. Prior to the induction of anesthesia, a premedication of 0.2 mg/kg atropine (atropine-sulfate, Braun Melsungen AG, Melsungen) was given as an intraperitoneal (i.p.) injection to stabilize circulation. Anesthesia was induced in a plastic chamber, which was flooded with 5% Isoflurane (Florene®, AbbVie Deutschland GmbH & Co. KG, Ludwigshafen, Germany) in pure oxygen (0.4 l/min). For maintenance of anesthesia, isoflurane was delivered through a breathing mask with a flow rate of 0.35 l/min and its concentration was regulated to 1.5-2.2% using an adjustable vaporizer (Uno, Zevenaar, Netherlands). The depth of anesthesia was controlled by testing the toe pinch reflex and the breathing rate (1-2Hz). Body temperature (36°C) and respiration were constantly monitored. Lidocaine/prilocaine ointment (25 mg/g, Emla® creme, AstraZeneca GmbH, 22876 Wedel) was applied prior to surgery and after suturing of the wound as local anesthetics. Additional analgesia was provided by subcutaneous injection of carprofen (4 mg/kg in NaCl, Rimadyl®, Pfilzer GmbH, Berlin, Germany) after infusion. Eye lubricant (Visidic, Bausch and Lomb, Berlin, Germany) was used to protect eyes from desiccation.

Desipramine hydrochloride (20 mg/kg, Sigma Aldrich) was injected i.p. 20-40 min before intracranial infusions to prevent 6-OHDA uptake by noradrenergic neurons. The desipramine solution was prepared in sterile, isotonic NaCl solution (B. Braun Melsungen AG, Germany) at the day of surgery. The infusion solutions are based on sterile artificial cerebrospinal fluid (ACSF, Harvard Apparatus, Holliston, MA, USA) with 0.02% L-ascorbic acid (used also as a vehicle solution). The 6-OHDA solution (0.2% 6-hydroxydopamine hydrochloride in ACSF with 0.02% L-ascorbic acid) was prepared at the day of infusion, stored on ice, and shielded from light.

During surgery, the animals were placed on a heating pad and were fixed in a stereotactic frame (Model 1900, Kopf Instruments, Tujunga, USA) with a stereotactic arm and a connected three-way digital positioning display. The scalp was opened by a longitudinal cut to expose the skull with bregma and lambda on display. With a centering scope (Model 1915, Kopf Instruments, Tujunga, USA), the bregma-lambda distance was measured and examined for suitable anatomy (4.4 ± 0.2 mm distance). Afterwards, the skull was aligned to a reference frame with a stereotaxic alignment indicator (Model 1905, Kopf Instruments, Tujunga, USA) and the manipulator system was referenced to bregma.

Using a stereotaxic drill (Model 1911, Kopf Instruments, Tujunga, USA) with a 500 µm diameter drill bit, a hole above the right striatum was drilled (coordinates: ML: +1.9 mm, AP: +0.5 mm to bregma). ACSF or 6-OHDA solution were loaded to a 10 µl NanoFil syringe (World Precision Instruments Inc., Sarasota, FL, USA) with a 35G blunt needle, which was mounted on a MicroSyringe Pump (UMP3-1, World Precision Instruments) and controlled by a SYSMicro4 Controller (World Precision Instruments). Using the stereotactic arm, the needle was slowly lowered (about 750 µm/min) to a position of –2.2 mm below the brain surface (infusion site coordinates: ML: +1.9 mm, AP: +0.5 mm, DV: –2.2 mm to bregma). Anatomical references are based on Franklin and Paxinos (2008). A volume of 6 µl was infused with a flow rate of 250 nl/min. Once the volume was infused, the needle rested for 5 minutes in that position before it was slowly moved out of the brain. Directly before and after infusion, proper functioning of the syringe system and the needle was checked. Finally, after suture the animal was placed on a heating pad for full recovery. Oats, wet food pallets and water were placed inside the cage to ease consumption.

### Behavioral testing

#### Open field

Spontaneous locomotion (track length, wall distance, time in center and number of rearings) and rotations of all mice were monitored in open field (50 × 50 cm, center 30 × 30 cm; red illumination, 3 lx) for 10 min in 3 baseline sessions and every 4^th^ or 7^th^ day post-infusion of ACSF/6-OHDA till the day of *in vivo* or *in vitro* experiment (e.g. 21^st^ or >68^th^ post-operative day). The open field was cleaned before and after each mouse with 0,1% acetic acid in distilled water. Using a video tracking system (Viewer II/III, Biobserve) spontaneous behavior was recorded and analyzed both online and offline. Data was extracted from Viewer as Excel-tables and the final analysis was made with custom made Matlab-scripts.

#### Cylinder test

Forelimb use during explorative activity was explored with cylinder test. The test was performed at corresponding termination time point (20-21^st^ and 64^th^ post-infusion day). Mice were placed individually in a glass beaker (9 cm diameter, 19 cm height) at room light and were video recorded with a camera (Logitech HD Webcam C615) for about 5 min. No habituation was allowed before video recording. The glass cylinder was cleaned before and after every mouse with 0,1% acetic acid in distilled water. Only weight-bearing wall contacts made by each and both forelimb on the cylinder wall were scored. Wall exploration was expressed in terms of the percentage of contralateral to the infusion side (in the 6-OHDA-infused mice also impaired forepaw) to all forelimb wall contacts.

### *In vivo* electrophysiology

#### Extracellular recording

*In vivo* extracellular single-unit activities of SN and VTA neurons were recorded in ACSF-infused (vehicle) and 6-OHDA-infused mice, similar procedures were used in other studies from our lab (Farassat et al. 2019; Schiemann et al. 2012; Subramaniam, Althof, et al. 2014). Briefly, mice were anesthetized (isoflurane; induction 4.5-5%, maintenance 1–2% in 0.4 l/min O2) and placed into a stereotactic frame. The craniotomies were performed as described above to target the lateral SN (AP: –3.08 mm, ML: 1.4 mm) and medial SN (AP: –3.08 mm, ML: 0.9 mm). Borosilicate glass electrodes (10– 25 MΩ; Harvard Apparatus, Holliston, MA, USA) were made using a horizontal puller (DMZ-Universal Puller, Zeitz, Germany) and filled with 0.5 M NaCl, 10 mM HEPES (pH 7.4) and 1.5% neurobiotin (Vector Laboratories, Burlingame, CA, USA). A micromanipulator (SM-6; Luigs & Neumann, Ratingen, Germany) was used to lower the electrodes to the recording site. The single-unit activity of each neuron was recorded for at least 10 minutes at a sampling rate of 12.5 kHz (for firing pattern analyses), and then for another one minute at a sampling rate of 20 kHz (for the fine analysis of AP waveforms). Signals were amplified 1000x (ELC-03M; NPI electronics, Tamm, Germany), notch-and bandpass-filtered 0.3–5000 Hz (single-pole, 6 dB/octave, DPA-2FS, NPI electronics) and recorded on a computer with an EPC-10 A/D converter (Heka, Lambrecht, Germany). Simultaneously, the signals were displayed on an analog oscilloscope and an audio monitor (HAMEG Instruments CombiScope HM1508; AUDIS-03/12M NPI electronics). Midbrain DA neurons were initially identified by their broad biphasic AP (> 1.2 ms duration) and slow frequency (1–8 Hz) (Anthony A. Grace and Bunney 1984; Ungless and Grace 2012). AP duration was determined as the interval between the start of initial upward component and the minimum of following downward component.

#### Juxtacellular labeling of single neurons

In order to identify the anatomical location and neurochemical identity of the recorded neurons, they were labeled post-recording with neurobiotin using the juxtacellular *in vivo* labeling technique (Pinault 1996). Microiontophoretic currents were applied (1–10 nA positive current, 200ms on/off pulse, ELC-03M, NPI Electronics) via the recording electrode in parallel to the monitoring of single-unit activity. Labeling was considered successful, when: the firing pattern of the neuron was modulated during current injection (i.e., increased activity during on-pulse and absence of activity in the off-pulse), the process was stable for at least 20s, and was followed by the recovery of spontaneous activity. This procedure allowed for the exact mapping of the recorded DA neuron within the SN and VTA subnuclei (Franklin and Paxinos 2012) using custom written scripts in Matlab (MathWorks, Natick, MA, USA), combined with neurochemical identification using TH-immunostaining.

### *In vitro* electrophysiology

#### Slice preparation

Animals were anesthetized by intraperitoneal injection of ketamine (250 mg/kg, Ketaset, Zoetis) and medetomidine-hydrochloride (2.5 mg/kg, Domitor, OrionPharma) prior to intracardial perfusion using ice-cold ACSF consisting of the following (in mM): 50 sucrose, 125 NaCl, 2.5 KCl, 25 NaHCO3, 1.25 NaH2PO4, 2.5 glucose, 6 MgCl2, 0.1 CaCl2 and 2.96 kynurenic acid (Sigma-Aldrich), oxygenated with 95% O2 and 5% CO2. Rostral coronal midbrain slices (bregma: –2.92 mm to –3.16 mm) were sectioned at 250 µm using a vibrating blade microtome (VT1200s, Leica). Slices were incubated for 1 h before recordings in a 37°C bath with oxygenated extracellular solution with extra 1µM AmmTx3, containing the following (in mM): 22.5 sucrose, 125 NaCl, 3.5 KCl, 25 NaHCO3, 1.25 NaH2PO4, 2.5 glucose, 1.2 MgCl2 and 1.2 CaCl2.

#### In vitro patch-clamp recordings

Slices were placed in a heated recording chamber (37°C) that was perfused with oxygenated extracellular solution at 2-4 ml min_−1_. CNQX (20 μM; Biotrend), gabazine (SR95531, 4 μM; Biotrend) DL-AP5 (10 μM; Cayman Chemical) were added to inhibit excitatory and inhibitory synaptic transmission. For voltage clamp recordings, TTX (500nM; Tocris) was added to the extracellular solution. Neurons were visualized using infrared differential interference contrast videomicroscopy with a digital camera (VX55, Till Photonics) connected to an upright microscope (Axioskop 2, FSplus, Zeiss). Patch pipettes were pulled from borosilicate glass (GC150TF-10; Harvard Apparatus LTD) using a temperature-controlled, horizontal pipette puller (DMZ-Universal Puller, Zeitz). Patch pipettes (4-6 MΩ) were filled with a solution containing the following (in mM): 135 KGlu, KCl, 10 HEPES, 0.1 EGTA, 5 MgCl2, 0.075 CaCl2, 5 NaATP, 1 LiGTP, 0.1% neurobiotin, adjusted to a pH of 7.35 with KOH. Recordings were performed using an EPC-10 patch-clamp amplifier (Heka electronics) with a sampling rate of 20 kHz and a low-pass filter (Bessel, 5 kHz). For voltage clamp recordings only experiments with uncompensated series resistance <10MO were included in this study and series resistance was electronically compensated 75%. Neurons were held at a holding potential of –40mV to minimize HCN activation. To determine Kv4.3 activation kinetics neurons were hyperpolarized to –80mV for 500ms followed by varying voltage steps from –60mV to –20mV in increments of 5mV for 1s. For inactivation kinetics neurons were hyperpolarized from –120mV to –20mV in increments of 10 mV for 1s followed by a fixed voltage step to –20mV for 1s. For analysis of HCN currents neurons were hyperpolarized from –80mV to –120mV in increments of 20mV for 1s. For analysis, action potential thresholds (mV) were determined at dV_m_/dt > 10 mV/ms.

### Immunohistochemistry

Following *in vivo* recordings, animals were transcardially perfused, as described previously (Farassat et al. 2019; Schiemann et al. 2012; Subramaniam, Althof, et al. 2014). Fixed brains were sectioned into 60µm (midbrain) or 100µm (forebrain) coronal sections using a vibrating microtome (VT1000S, Leica). *In vitro* slices were fixed in paraformaldehyde after finishing the experiment. Sections were rinsed in PBS and then incubated (in blocking solution (0.2 M PBS with 10% horse serum, 0.5% Triton X-100, 0.2% BSA). Afterwards, sections were incubated in carrier solution (room temperature, overnight) with the following primary antibodies: polyclonal guinea pig anti-tyrosine hydroxylase (TH; 1:1000; Synaptic Systems), monoclonal mouse anti-TH (1:1000; Synaptic Systems) or polyclonal rabbit anti-TH (1:1000; Synaptic Systems); mouse anti-Kv4.3 (1:1000, Alomone Labs). In sequence, sections were again washed in PBS and incubated (room temperature, overnight) with the following secondary antibodies: goat anti-guinea pig 488, goat anti-rabbit 488, goat anti-mouse 488, goat anti-mouse 568 (all 1:750; Thermofisher). Streptavidin AlexaFluor-568 and Streptavidin AlexaFluor-488 (both 1:1000; Invitrogen) were used for identifying neurobiotin-filled cells. Sections were then rinsed in PBS and mounted on slides with fluorescent mounting medium (Vectashield, Vector Laboratories).

### DAB immunocytochemistry

For DAB (3,3’-diaminobenzidine) staining procedures, a Vectastain ABC Staining Kit (Vector Laboratories) was used. Coronal sections of midbrain (30 μm) areas were cut and rinsed in PBS (3×10 min). Similar to previous immunolabeling procedures, unspecific antigen binding sites were blocked by incubation of the sections with blocking solution (60 min, room temperature). Subsequently, sections were incubated with primary antibody against TH (rabbit anti TH) overnight, rinsed in PBS (3×10 min), and were incubated with biotinylated secondary antibodies (biotinylated anti-rabbit) for two hours at RT. In parallel, an avidin-biotin complex (ABC) was formed by pre-incubation of avidin (1:1000) with biotinylated HRP (1:1000) in PBS for two hours at room temperature. Sections were rinsed in PBS (3×10 min) prior to incubation with ABC solution (60 min, room temperature). Next, sections were rinsed in PBS (2×10 min) and Tris-buffer (1×10 min). Finally, DAB oxidation was developed by application of 2 % H_2_O_2_, 2 % NiCl_2_ and 4 % DAB in Tris-buffer using a DAB Substrate Kit (Vector Laboratories, Burlingame, USA). NiCl_2_ enhances sensitivity and intensity of DAB precipitation product. DAB oxidation was developed for 2 to 5 minutes and was stopped with Tris-buffer once a specific high-contrast signal was detectable. Sections were rinsed in Tris-buffer (3×10 min) and transferred onto gelatin-covered slides, air-dried overnight, and dehydrated in consecutive ascending alcohol concentrations (50 %, 70 %, 90 % and 2× 100 %; 10 min each) followed by dehydration in xylol (2× 100 %; 10 min each). Finally, sections were mounted under glass coverslips with hardening mounting medium (Vectamount, Vector Laboratories, Burlingame, USA).

### Unbiased stereology measurements

For quantification of total cell loss, TH-DAB labeled SN DA neurons were counted using unbiased stereology based on optical dissection (Gundersen 1986). In coronal sections (30 μm), the region of interest was selected based on anatomical landmarks including the medial lemniscus, which separates SN and adjacent VTA. Stereological counting provides unbiased data based on random, systematic sampling using an optical fractionator. This method involves counting of neurons with an optical dissector, a three-dimensional probe placed through a reference space (Gundersen 1986). The optical dissector forms a focal plane with a thin depth of field through the selected sections. Objects in focus of this focal plane are located within the reference section and are counted, while objects outside of the focal plane are not counted. On top of the optical dissector, a counting frame is applied. Counting frames ensure that all neurons have equal probabilities of being selected, regardless of shape, size, orientation, and distribution. To avoid counting capped neurons at the border of a section, an additional guard zone was deployed at the upper and lower borders of each section. DA neurons within the counting frame as well as those crossing the green line (acceptance line) were counted, while DA neurons crossing the red line (rejection line) excluded. Moreover, only neurons with a detectable nucleus in focus within the optical dissector were counted. For quantification of total cell loss, StereoInvestigator software (V5, MicroBrightField, Colchester, USA) was used in combination with BX61 microscope (Olympus, Hamburg, Germany). The region of interest was selected and marked using a low magnification objective lens (2x, NA 0.25, Olympus) and 12-30 serial sections of 30 μm thickness were counted, covering the entire caudo-rostral extent of the SN. To count the number of DA neurons in the area of the SN pars compacta, a high magnification oil-immersion objective lens (100x, NA 1.30, Olympus) (counting frame, 50 × 50 μm; sampling grid, 125 × 100 μm) was used. After counting was finished, the total number of neurons was calculated by the software using the formula described by West et al (West 1993).

### Optical density measurements

Optical density measurements of TH-DAB labeled striatal sections were performed using ImageJ software (http://rsbweb.nih.gov/ij/). Following TH-DAB labeling and TH-immunohistochemistry, images of five coronal sections (100 μm) covering the rostrocaudal axis of the striatum were captured using laser-scanning microscope (Nikon Eclipse90i, Nikon GmbH). Images were gray scale converted and mean gray values of desired striatal areas were encircled and measured. Unspecific mean gray values were measured in a defined cortical area (100×100 pixels) that displayed no specific TH signal due to the absence of DA innervation and were subtracted. The ventral edge of lateral ventricles served as an anatomical landmark to separate dorsal and ventral areas. For all animals, the measurement from the ipsilateral to the infusion side were divided by the contralateral side to calculate the relative optical density of the striatum.

### Immunohistochemical Kv4.3 channel signal quantification

A Nikon Eclipse 90i microscope was used for fluorescent signal detection, excitation wavelength of 488 nm for TH-signal and 568 nm for Kv4.3-channel signal. From each animal, 4 midbrain slices covering the caudal, intermedial and rostral regions, were selected and imaged for overview with 4x magnification. Then 60x magnification was used to acquire data from 4 areas within each SN (4 images on the ipsilateral and 4 images on the contralateral to infusion side). All images were acquired using the same laser and camera settings. Images were exported from Nikon NIS-Elements Advanced Research (Version 4.20.03) software as 8-bit TIFF files for quantification. Data was analyzed using custom made Python 3.0 scripts with matplotlib, numpy, scimage and scipy modules. First, TH immunosignals were converted to a binary image via Otsu-thresholding algorithm to detect TH areas bigger than 400 pixels. Then, the resulting binary image was used as a mask for Kv4.3 channel immunosignal detection. For all ROIs surface areas and mean Kv.4.3 channel signal intensity was measured. By applying erosion and dilatation algorithms on the ROIs, membrane and cytoplasm areas were segregated, allowing isolation of Kv4.3 channel immunosignal intensity for these cell compartments. Background Kv4.3 channel immunosignal was quantified in TH-immunosignal areas below the Otsu-threshold. All data were then grouped according to medio-lateral and ipsi-/contralateral position for both vehicle and 6-OHDA group. Graphs and statistical analysis for this data was performed using Python custom made scripts.

### Statistical Analysis

#### Spike train analyses

Spike time-stamps were extracted by thresholding above noise levels with IgorPro 6.02 (WaveMetrics, Lake Oswego, OR, USA). Firing pattern properties such as mean frequency, coefficient of variation (CV) and bursting measures were analyzed using custom scripts in Matlab. In order to estimate burstiness and intra-burst properties, we used the burst detection methods described in Grace & Bunney (A. A. Grace and Bunney 1984; Anthony A. Grace and Bunney 1984; Ungless and Grace 2012). All non-burst related ISIs (excluding all ISIs that followed the Grace and Bunney criteria, as well as all pre– and post-burst ISIs) were used to calculate the single spike firing frequency and single spike coefficient of variation.

For analysis of general firing patterns, autocorrelation histograms (ACH) were plotted using custom Matlab scripts. We used established criteria for classification of *in vivo* firing patterns based on visual inspection of autocorrelograms (Farassat et al. 2019; Schiemann et al. 2012; Subramaniam, Althof, et al. 2014): single spike-oscillatory (≥3 equidistant peaks with decreasing amplitudes), single spike-irregular (<3 peaks, increasing from zero approximating a steady state), bursty-irregular (narrow peak with steep increase at short ISIs) and bursty-oscillatory (narrow peak reflecting fast intraburst ISIs followed by clear trough and repetitive broader peaks).

#### Statistics

Categorical data is represented as stacked bar graphs. To investigate the assumption of normal distribution, we performed the single-sample Kolmogorov-Smirnov test. The Mann-Whitney-Test, one-/two-way ANOVA were performed in non-parametric data to determine statistical significance. Categorical parameters, such as ACH-based firing pattern, were analyzed with the Chi-squared test. Statistical significance level was set to *p* < 0.05. All data values are presented as means ± SEM. Statistical tests were performed using GraphPad Prism 9 (GraphPad Software, San Diego, CA, USA), Matlab and Python. The scatter plots are represented with median or mean ± SEM. The resulting *p* values were compared with Bonferroni-corrected α-level or Tukey *post hoc* comparison. A value of *p* ≤ 0.05 was considered to be statistically significant; *p* ≤ 0.05 = \**p* ≤ 0.005 = \*\**p* ≤ 0.0005 = ***. Graphs were plotted using GraphPad Prism software (9.0c), Matlab and Python.

## Acknowledgements

This study was supported by research grants to JR (DFG, CRC 1451). LK is a MD/PhD candidate at TransMed, Gutenberg University Mainz. We thank Beatrice Fischer and Jasmine Sonntag for technical assistance, Alexander Prinz for preliminary data on post-6-OHDA stereology.

## Additional information Author contributions

LK & JR designed the study. LK performed the lesions, behavioral & *in vivo* electrophysiology, JS & JM performed the *in vitro* experiments. JZ performed dose response lesion and female experiments. Analysis was carried out jointly by LK, JZ, KMC, JO and JR. NF taught LK *in vivo* electrophysiology and juxtacellular labelling. LK & JR wrote the manuscript.

## Ethics

Animal experimentation: All experiments and procedures involving mice were approved by the German Regierungspräsidium Darmstadt (V54-19c20/15-F40/30).

## Supplementary Figures

**Supplementary Figure 1.**
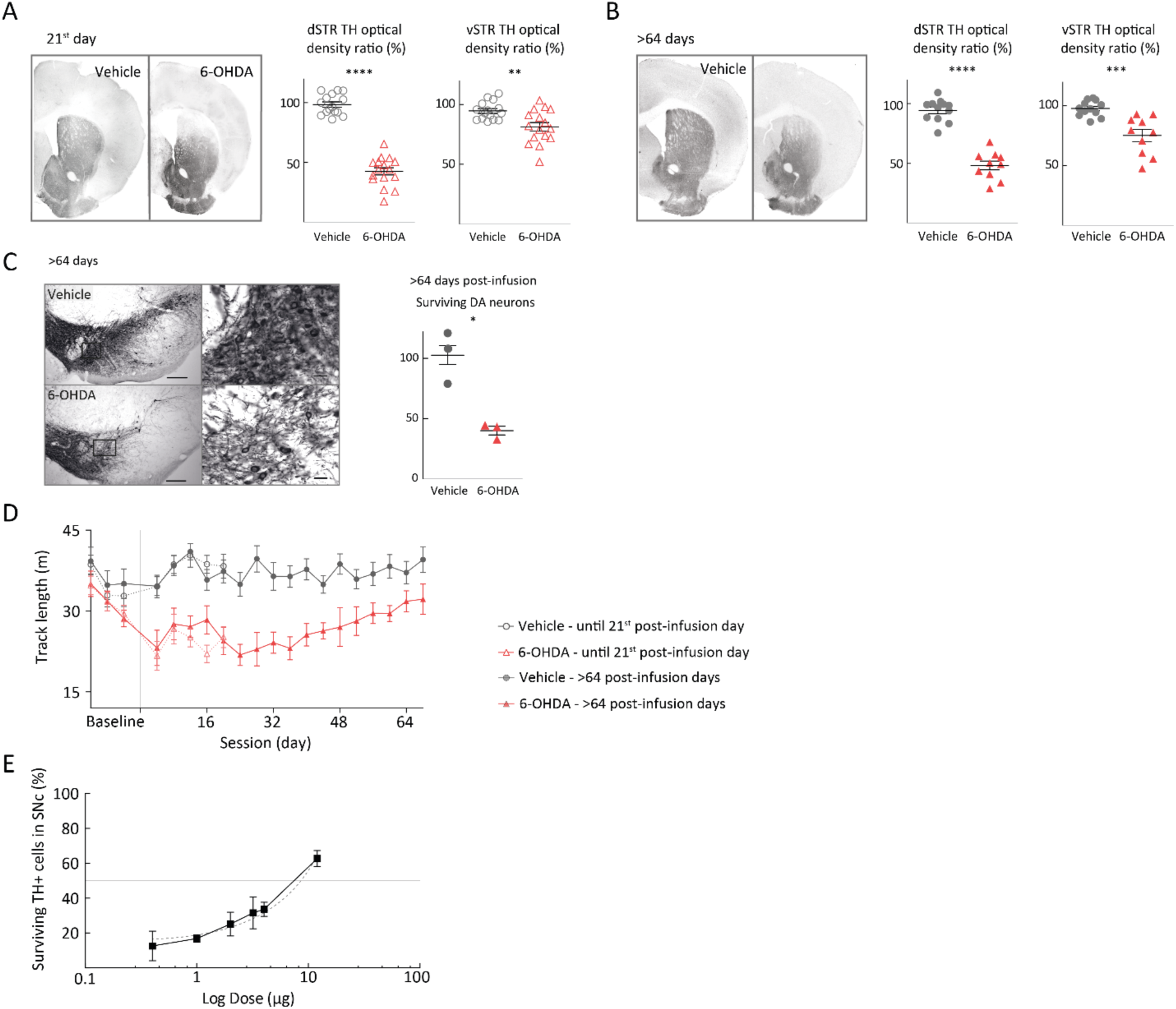
(A), (B) Comparison of ipsilateral, infusion side, to contralateral side as percentage of relative TH immunohistochemistry signal in the dorsal striatum (dSTR) and ventral striatum (vSTR), at the corresponding early (A) and late phase (B). Note the stability of TH density loss in the dorsal striatum through time. (C) Ratio of ipsilateral (infusion side) to contralateral side of surviving TH-positive neurons in SN in the late phase. (D) Mean track length from all mice for each open field session. Note the post-infusion drop in performed track in the 6-OHDA groups, which gradually recovers. (Infusion day marked as a thin gray line.) (E) Ratio of TH+ cell loss in SNc (ipsilateral (infusion side) to contralateral side) >21st post-infusion day from different infused 6-OHDA doses.

**Supplementary Figure 2.**
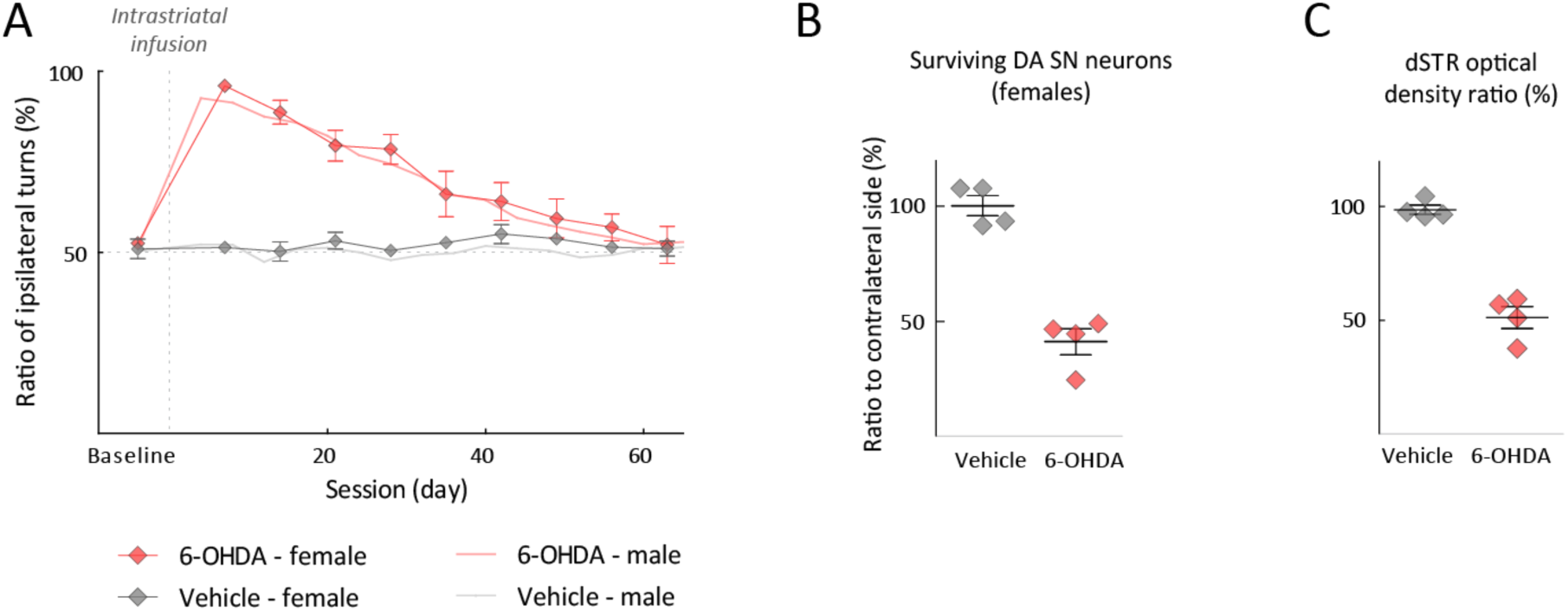
(A) Ratio of ipsilateral to contralateral turning behavior for female ACSF and 6-OHDA infused groups plotted against session days and with underlayed the corresponding ratio from the male ACSF and 6-OHDA groups as a mean-line. (B) Ratio of ipsilateral (infusion side) to contralateral side of surviving TH-positive neurons in SN at >64st post-infusion day from female ACSF and 6-OHDA infused mice. (C) Comparison of ipsilateral, infusion side, to contralateral side as percentage of relative TH immunohistochemistry signal in the dorsal striatum (dSTR) at >64st post-infusion day from female ACSF and 6-OHDA infused mice.

**Supplementary Figure 3.**
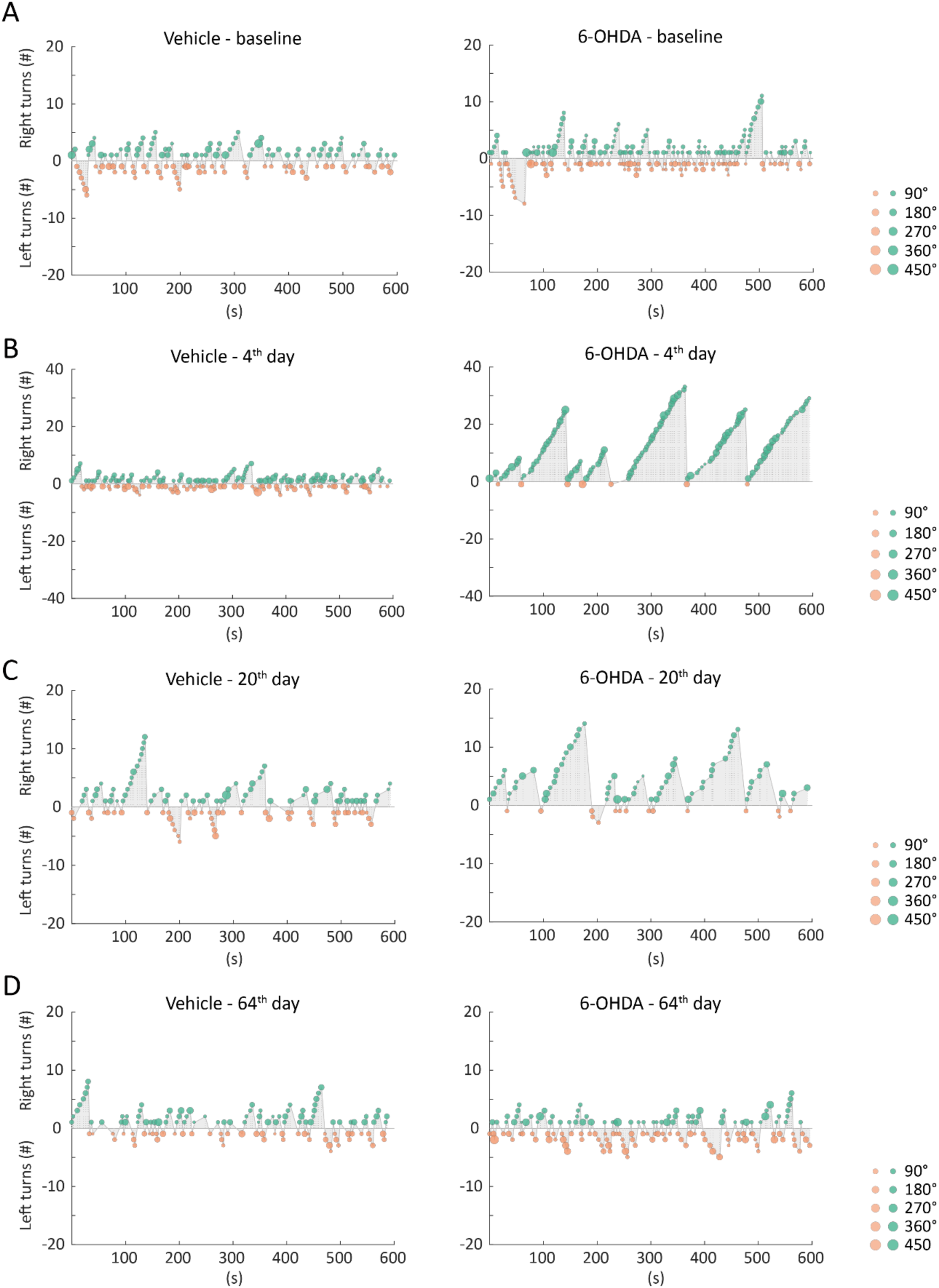
For two representative vehicle– and 6-OHDA-treated mice schematic representation of turning sequences. Left turn sequences are plotted as negative values, whereas right turn sequences are plotted as positive values. The count represents the number of same-direction turns within a sequence. On the left panels are turning sequences for the control animal and on the right panels for the 6-OHDA-infused animal, correspondingly in a baseline session (A), 4^th^ post-operative day (B), 20^th^ post-operative day (C) and 64^th^ post-operative day (D).

**Supplementary Figure 4.**
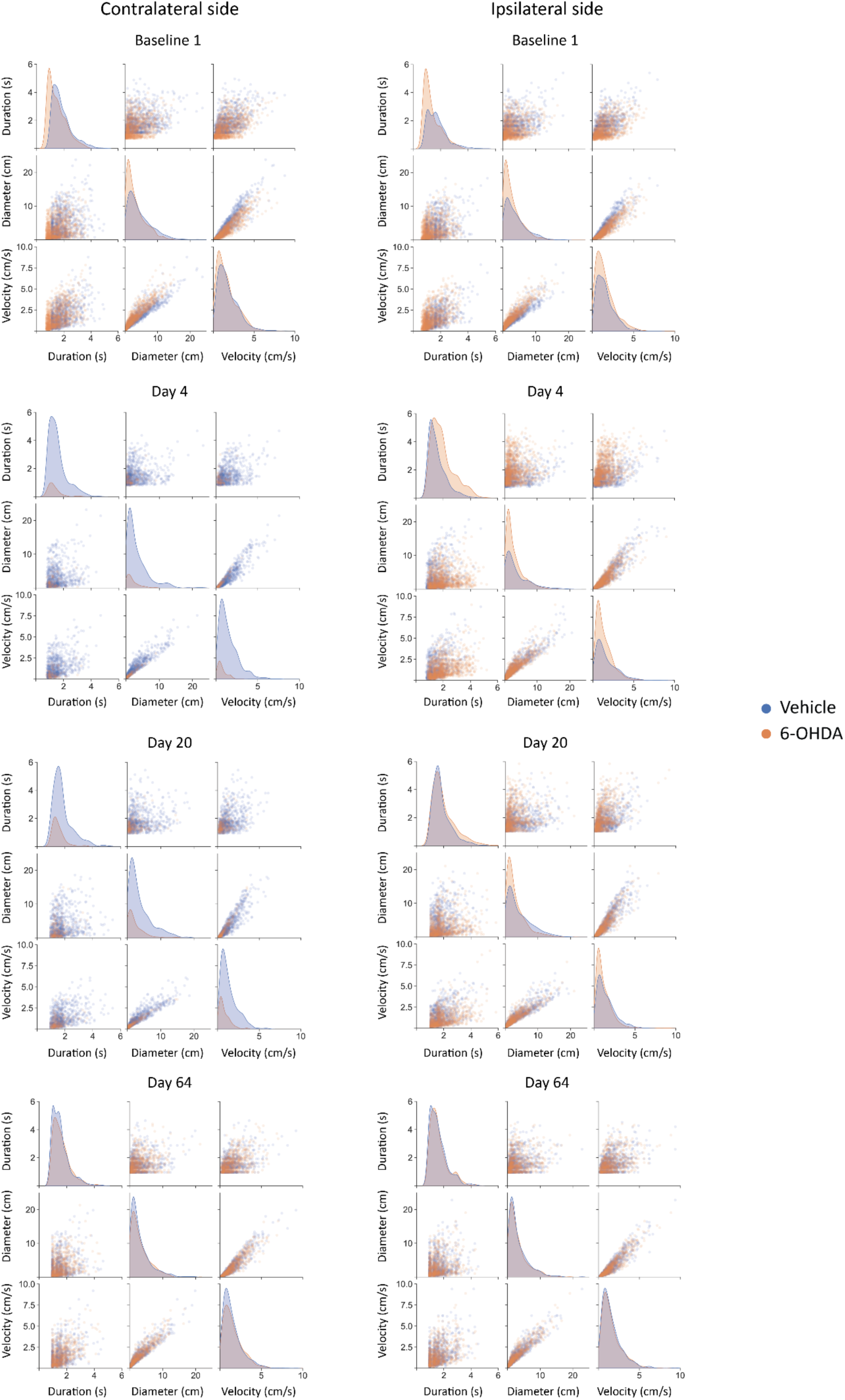
Correlation pair plots comparing turning features, such as velocity (cm/s), diameter (cm) and duration for all contralateral turns (left panels) and all ipsilateral turns performed from all vehicle-infused mice (in orange) and all 6-OHDA-treated mice (in blue) across different time points (baseline, 4^th^, 20^th^ and 64^th^ post-infusion day).

**Supplementary Figure 5.**
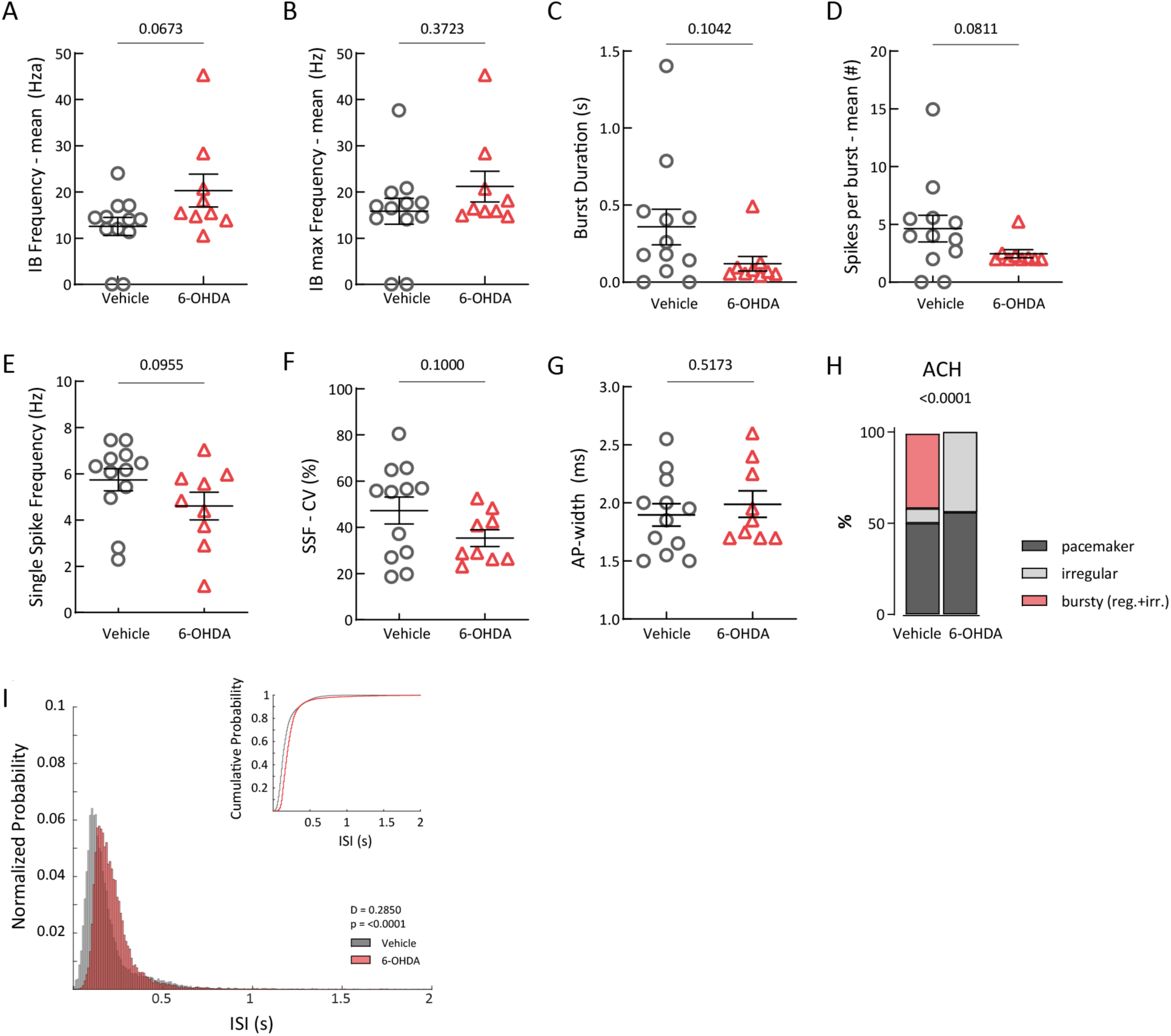
Scatter dot-plots, showing no significant difference of in vivo mean intra-burst (IB) frequency (A), mean maximum (max) intra-burst (IB) frequency (B), burst duration (C), number of spikes per burst (D), single spike frequency (SSF) (E), single spike coefficient of variance (CV) (F) and action potential (AP) width (G) (H) between the vehicle and 6-OHDA-infused mice in the early phase. (H) Normalized stacked bar plots of different in vivo firing patterns based on ACH. (I) ISI distributions from all in vivo recorded and labeled mSN DA neurons from vehicle– and 6-OHDA-treated mice in the early phase. Inset, cumulative representation of the same distributions.

**Supplementary Figure 6.**
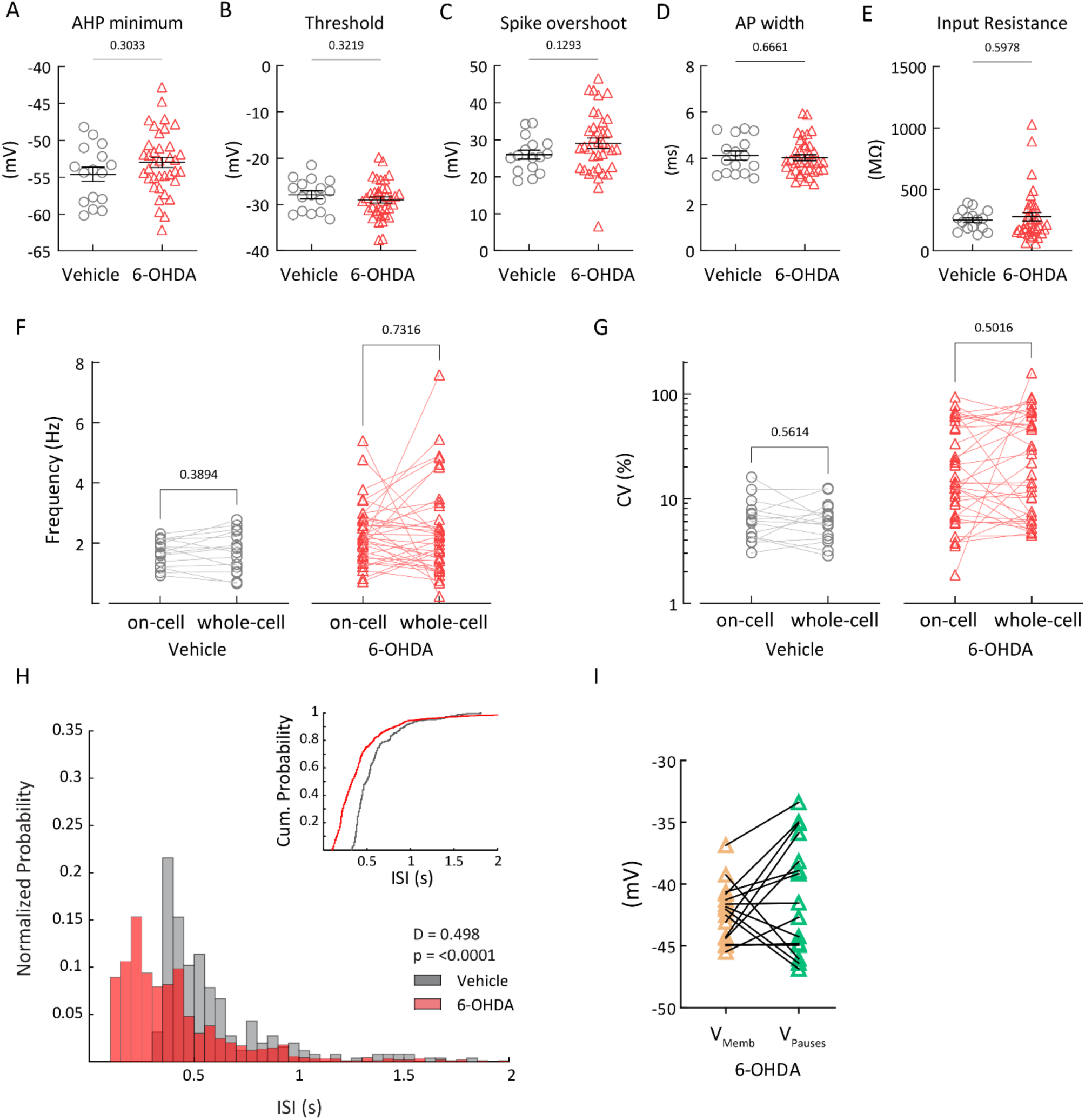
Scatter dot-plots, showing no significant difference of in vitro afterhyperpolarization (AHP) minimum (A), threshold (B), spike amplitude (C), action potential (AP) width (D) and input resistance (E) between the vehicle and 6-OHDA-infused mice in the early phase. (F), (G) Paired scatter dot-plots of firing frequency (F) and coefficient of variance (G) during on-cell recording and whole-cell recording in the early phase. (H) ISI distributions from all in vitro whole-cell recorded and labeled mSN DA neurons from vehicle– and 6-OHDA-treated mice in the early phase. Inset, cumulative representation of the same distributions. (I) Paired scatter dot-plot indicating no significant difference between the mean voltage during spike-pauses and the rest of the firing activity.

**Supplementary Figure 7.**
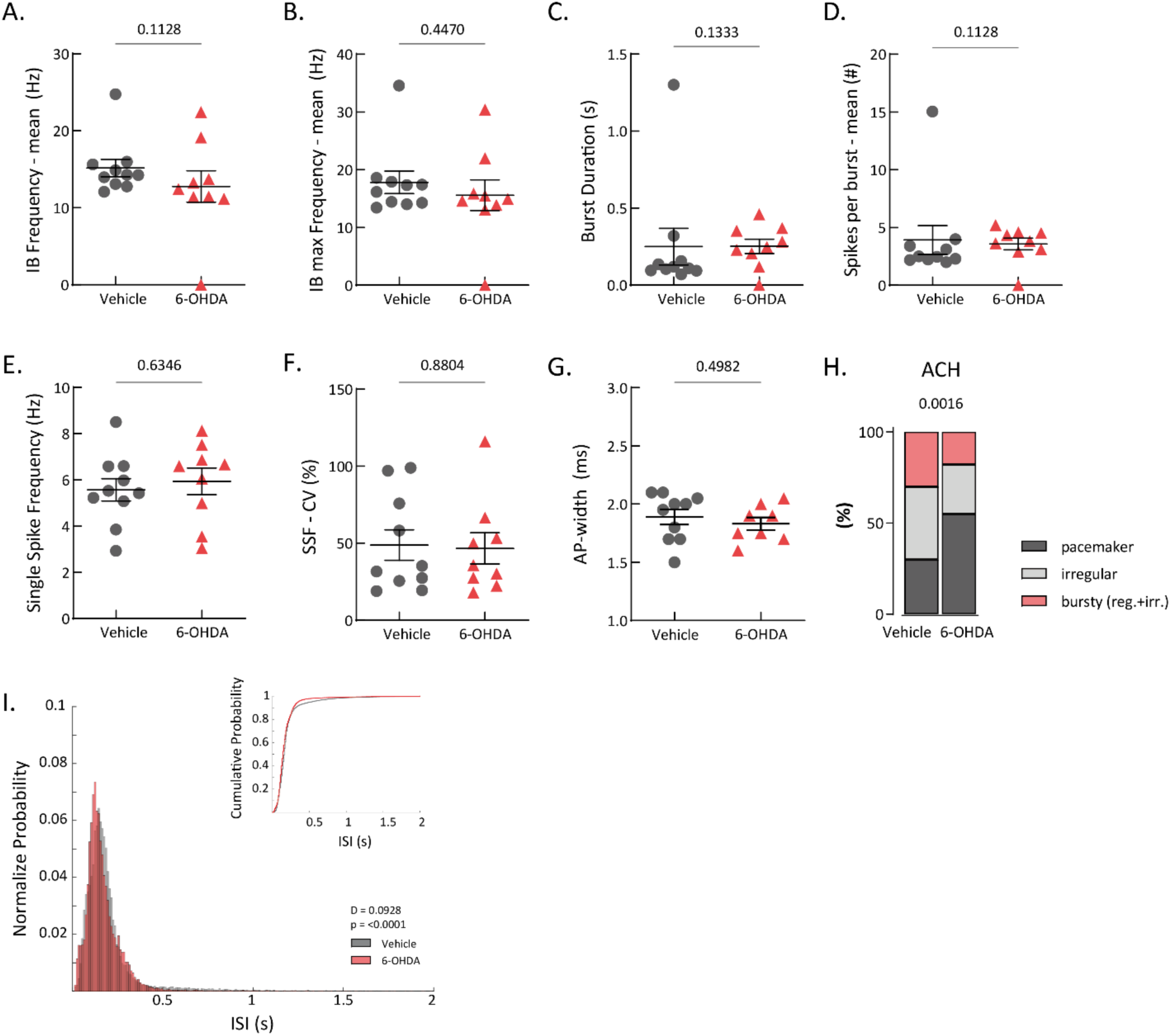
Scatter dot-plots, showing no significant difference of in vivo mean intra-burst (IB) frequency (A), mean maximum (max) intra-burst (IB) frequency (B), burst duration (C), number of spikes per burst (D), single spike frequency (SSF) (E), single spike coefficient of variance (CV) (F) and action potential (AP) width (G) (H) between the vehicle and 6-OHDA-infused mice in the late phase. (H) Normalized stacked bar plots of different in vivo firing patterns based on ACH. (I) ISI distributions from all in vivo recorded and labeled mSN DA neurons from vehicle– and 6-OHDA-treated mice in the late phase. Inset, cumulative representation of the same distributions.

**Supplementary Figure 8.**
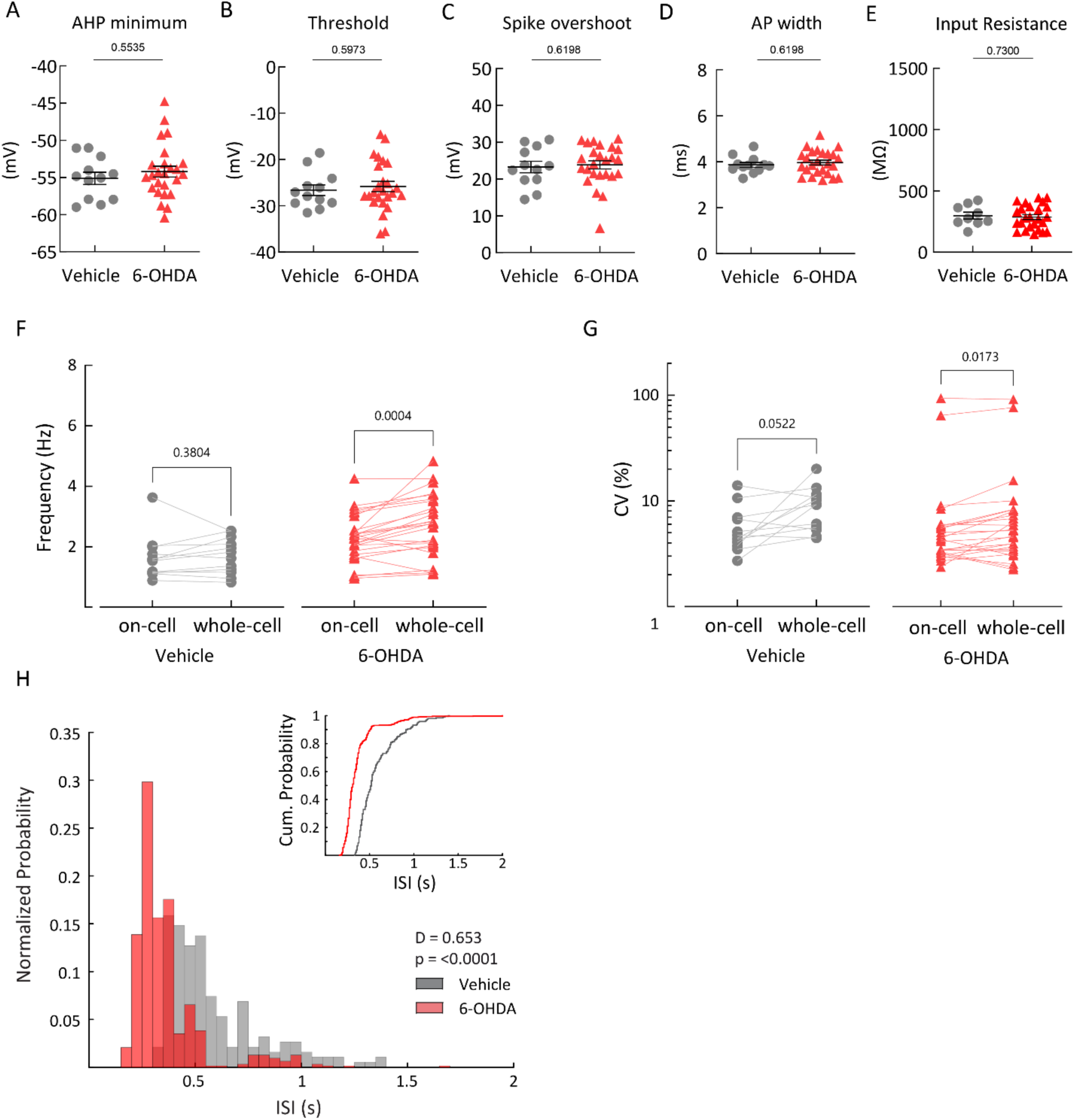
Scatter dot-plots, showing no significant difference of in vitro afterhyperpolarization (AHP) minimum (A), threshold (B), spike amplitude (C), action potential (AP) width (D) and input resistance (E) between the vehicle and 6-OHDA-infused mice in the late phase. (F), (G) Paired scatter dot-plots of firing frequency (F) and coefficient of variance (G) during on-cell recording and whole-cell recording in the late phase. (H) ISI distributions from all in vitro whole-cell recorded and labeled mSN DA neurons from vehicle– and 6-OHDA-treated mice in the late phase. Inset, cumulative representation of the same distributions.

**Supplementary Figure 9.**
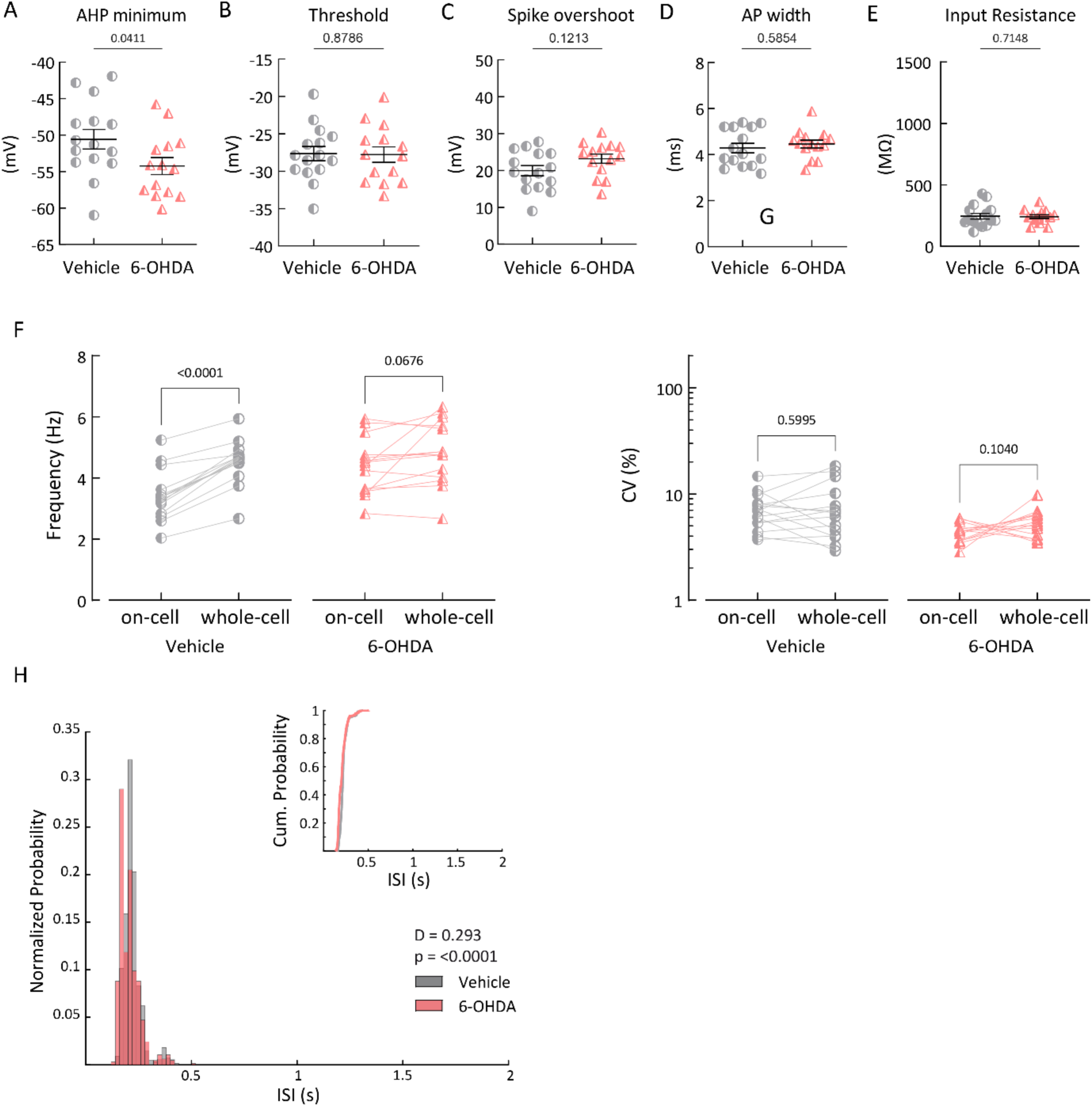
Scatter dot-plots, showing minor significant difference of in vitro afterhyperpolarization (AHP) minimum (A), and no difference in threshold (B), spike amplitude (C), action potential (AP) width (D) and input resistance (E) between the vehicle and 6-OHDA-infused mice in the late phase under 1µM AmmTx3. (F), (G) Paired scatter dot-plots of firing frequency (F) and coefficient of variance (G) during on-cell recording and whole-cell recording in the late phase under 1µM AmmTx3. (H) ISI distributions from all in vitro whole-cell recorded and labeled mSN DA neurons from vehicle– and 6-OHDA-treated mice in the late phase under 1µM AmmTx3. Inset, cumulative representation of the same distributions.

**Supplementary Figure 10.**
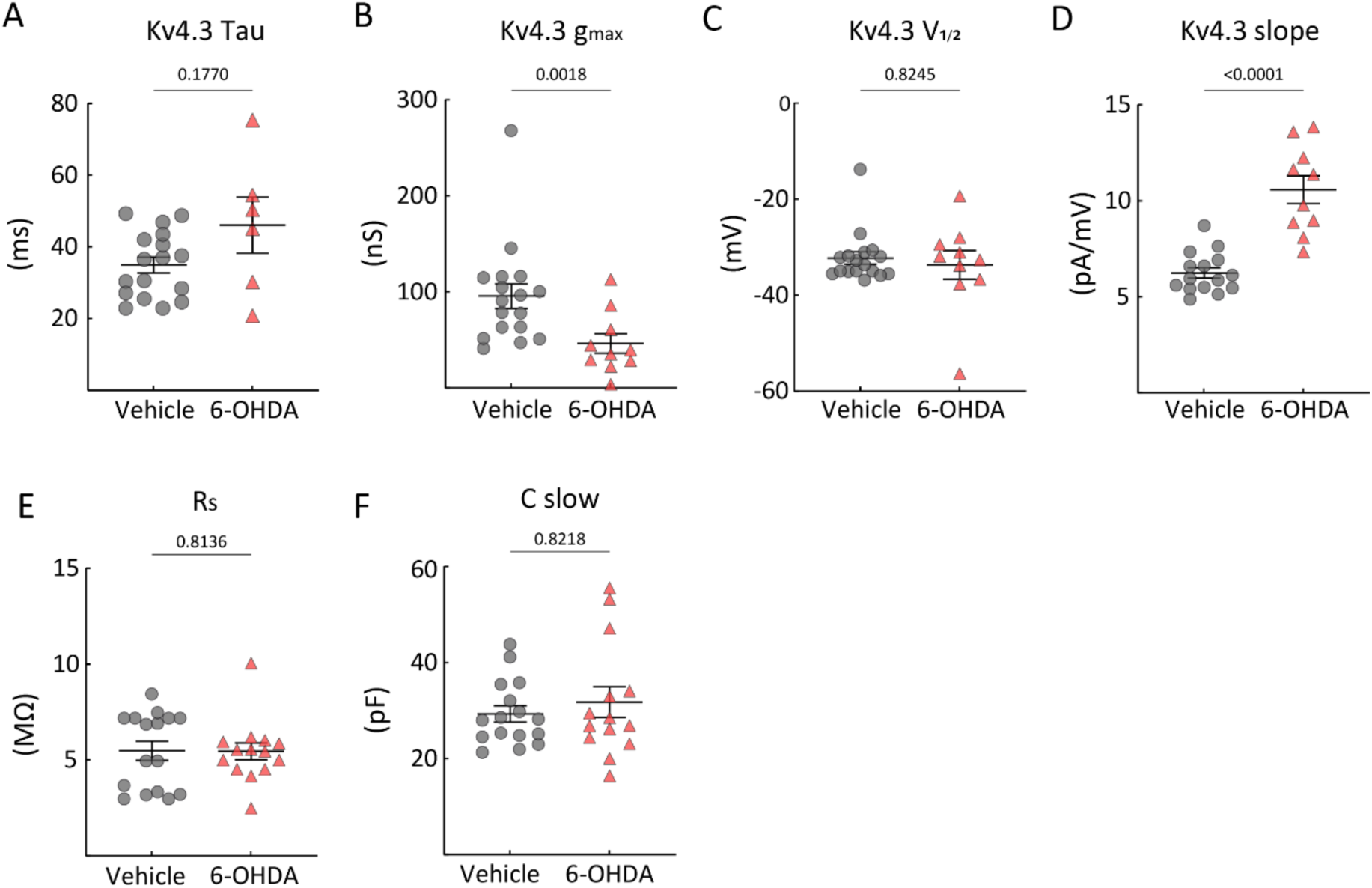
(A)-(D) Measurements of transient outward (A-type) potassium biophysical parameters in whole-cell voltage-clamp recordings of Kv4.3 channel activation and inactivation in vehicle relative to 6-OHDA infused mice. (E), (F) No difference in series resistance (Rs) or in slow capacitance between groups.

**Supplementary Figure 11.**
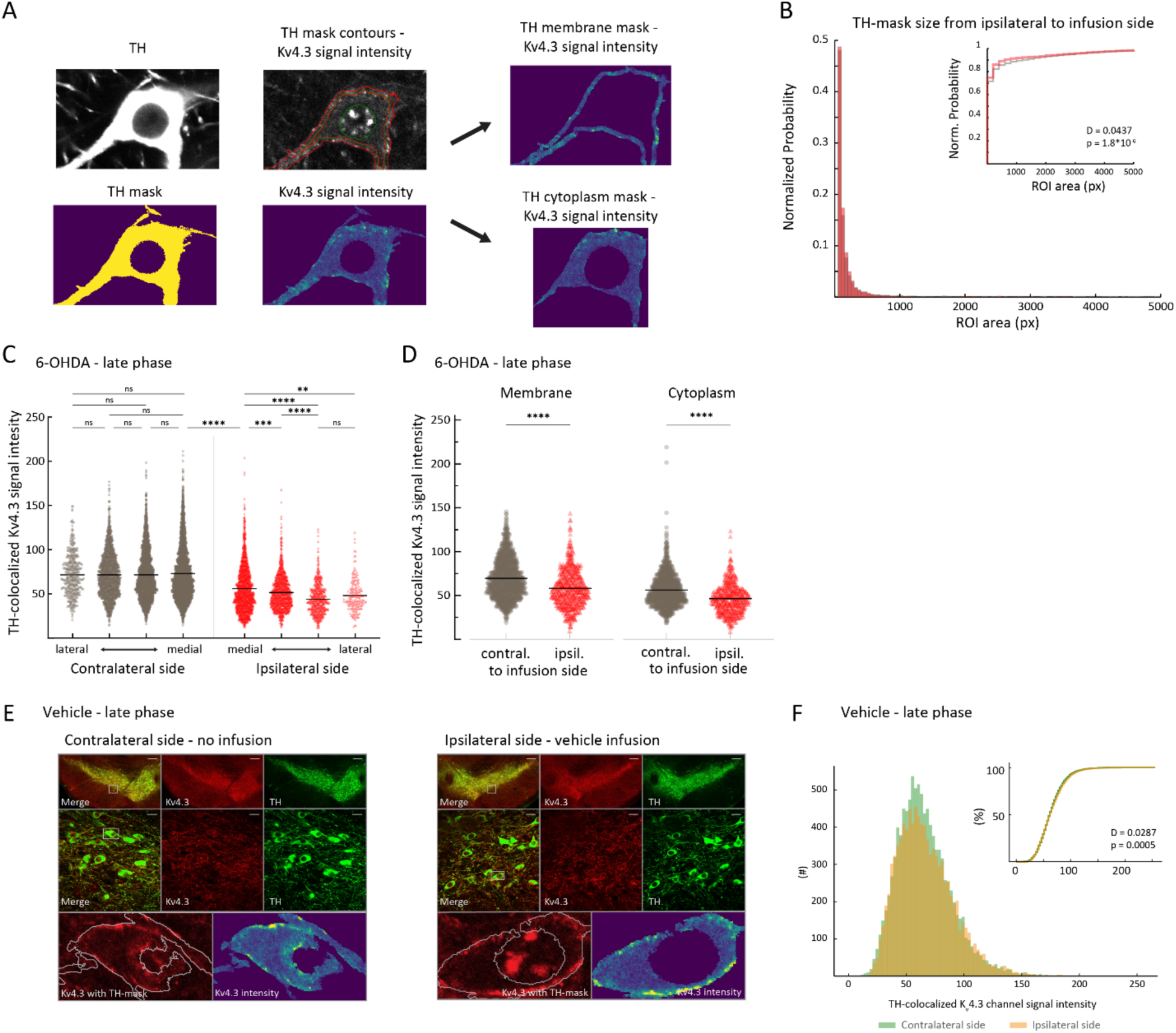
(A) Example illustration of an image from neuronal TH-immunosignal, transformed as a TH mask and overlayed on top of Kv4.3 immunosignal. Further segregation of the membrane and cytoplasm compartment with the corresponding Kv4.3 immunosignal intensity. (B) Distributions of all ROIs area sizes based on TH-masks from vehicle-(gray) and 6-OHDA-treated mice (red) in the late phase. Inset, cumulative representation of the same distributions. (C) Scatter dot-plots, illustrating the medio-lateral gradient of Kv4.3 immunosignal on the contralateral and ipsilateral side from 6-OHDA-treated mice in the late phase. (D) Scatter dot-plots, showing significant decrease in Kv4.3 immunosignal in both membrane and cytoplasm compartments in the ipsilateral (6-OHDA-infused) side compared to the contralateral (control) side. Membrane: p = 0.0003, contralateral side 67.2 ± 0.9, ipsilateral side 56.7 ± 1.1 (about 16% reduction); Cytoplasm: p < 0.0001, contralateral side 55.8 ± 0.9, ipsilateral side 45.9 ± 1.1 (about 18% reduction) (E) Top: 4x magnification of midbrain of a vehicle-infused mouse, >64 days post-lesion – contralateral side (left panel), and corresponding ipsilateral side (right panel). Middle: 60x magnification in the highlighted area from 4x image (green, TH; red, Kv4.3). Bottom, left: zoom-in on an example ROI (highlighted in 60x image). Bottom, right: color-coded Kv4.3-channel immunohistochemical signal intensity in the example ROI. (F) Histogram showing intensity of Kv4.3 immunosignals for all TH-positive ROIs, from ipsilateral, vehicle-infused, side (in orange) and from contralateral side (in green). Inset, same data shown as a cumulative distribution.

